# Drug-induced nuclear cathepsin L (nCTSL) defines a targetable DNA damage response axis and PARP inhibitor sensitivity in ovarian cancer

**DOI:** 10.1101/2025.01.09.632284

**Authors:** Prabhu Thirusangu, Ling Jin, Shruti Rao, Upasana Ray, Anya Zhao, Julie Staub, Xinyan Wu, Xiaonan Hou, Jamison L VanBlaricom, Syed Mohamed Musheer Aalam, Sun-Hee Lee, Ann L Oberg, Nagarajan Kannan, John Weroha, Jeremy Chien, Scott H Kaufmann, Jamie N Bakkum-Gamez, Viji Shridhar

## Abstract

**Background:** Despite treatment advances, ovarian cancer (OC) remains the deadliest gynecological cancer, with a ∼70% mortality rate. OCs initially respond to platinum chemotherapy but often develop resistance. While PARP inhibitors improve relapse-free survival in responsive cases, they do not extend overall survival. This underscores the need for effective combination therapies beyond BRCA status. The lysosomal protein cathepsin L (CTSL) can traffic to the nucleus, where it can regulate DNA repair and serve as a chromatin modifier. Here, we tested the hypothesis that nucleoside analog clofarabine (CLF) induced nuclear CTSL (nCTSL) serves as a biomarker of sensitivity to DNA-damaging agents in ovarian cancer.

**Methods:** Role of nuclear CTSL in DNA damage, synergy with PARPis and CRM1 inhibitor, tumor regression was analyzed by 3D-spheroid culture, colony forming assay, subcellular fractions, comet assay, and IFC/ western-blotting using PARP inhibitor-sensitive and-resistant OC cell lines, ex vivo cultures of patient-derived OC ascites cells (OVA), primary OCs, and patient-derived xenografts (PDXs).

**Results:** In OC cell lines, treatment with CLF induced nCTSL in a subset of models-designated CLF-responsive (CLF-r)-and sensitized them to the PARP inhibitors olaparib and rucaparib. In CLF-non-responsive (CLF-nr) models, nCTSL trafficking and drug synergy were achieved with the CLF+olaparib combination. Synergy was observed in 47% of CLF-r and 24% of CLF-nr OVA samples. Mechanistically, CLF monotherapy induced nuclear import of CTSL via KPNB1 in CLF-r cells, while combination treatment with CLF+olaparib was required in CLF-nr cells. CLF also downregulated the nuclear export protein CRM1 in both CLF-r and CLF-nr models. CTSL knockdown conferred resistance to CLF+olaparib in both cohorts, supporting a functional role for nCTSL in therapeutic response. Notably, in a subset of OVAs (29%) classified as CLF-resistant (CLF-Res), CLF failed to downregulate CRM1; however, pharmacological inhibition of CRM1 with KPT8602 restored sensitivity to CLF+PARP inhibitor treatment. In vivo, CLF+olaparib treatment in CLF-r and CLF-nr PDX models led to enhanced DNA damage, reduced tumor burden, and prolonged survival.

**Conclusion:** Collectively, these findings identify nCTSL as a predictive marker of response to DNA-damaging therapies and support the CLF+olaparib combination as a promising strategy for overcoming drug resistance in OC via CTSL-mediated DNA damage.

## BACKGROUND

Despite advances in treatment, ovarian cancer remains the most lethal gynecological malignancy, with a mortality rate of approximately 70%. High-grade serous ovarian cancer (HGSOC), the most common and deadliest subtype of ovarian cancer, initially responds to platinum-containing chemotherapy but eventually acquires platinum resistance. Although PARP inhibitors have improved progression-free survival, their efficacy is limited by the absence of robust predictive biomarkers beyond BRCA mutation status and by the near-universal development of resistance [1, 2]. These limitations highlight a critical need to identify mechanisms that dynamically regulate DNA repair capacity and can be therapeutically exploited. Nuclear trafficking has emerged as an important but underexplored regulator of cellular signaling; however, whether drug-inducible nuclear trafficking events can directly reprogram the DNA damage response remains unknown. In this context, we investigated the role of cathepsin L (CTSL), a lysosomal protease previously reported to localize to the nucleus under specific conditions, as a potential mediator of therapy response.

CLF (2-chloro-9-[2′-deoxy-2′-fluoro-β-D-arabinofuranosyl]-9H-purine-6-amine; CLF-ara-A; CAFdA) (CLOLAR®), a nucleoside analog, has shown efficacy in recurrent hematological malignancies, including pediatric and adult acute lymphoblastic leukemia [3, 4], acute myeloid leukemia (AML) [5], and myelodysplastic syndrome (ClinicalTrials.gov Identifier: NCT0070001). In various Phase II studies, CLF monotherapy has shown an overall response rate of 46-55% and a complete response rate of approximately 40% in AML [5, 6]. CLF has also shown therapeutic efficacy alone and in combination with other drugs in a wide range of solid tumor cell lines and patient-derived xenografts (PDXs) [7–13] but has not been tested in ovarian cancer nor with PARP inhibitors. In Phase I clinical studies in adult solid tumor patients, CLF administered weekly for 3 weeks (days 1, 8, and 15) every 28 days was well tolerated and showed clinical activity, with the maximum tolerated dose estimated to be 128 mg/m² and plasma Cmax of 720 nM at that dose [14]. The reported anticancer activity of CLF involves inhibition of DNA synthesis, direct induction of apoptosis, and inhibition of ribonucleotide reductase, resulting in replication fork stalling and DNA damage [15].

As first reported by Goulet et al. [16], a unique isoform of the protease CTSL that can traffic to the nucleus (nCTSL) can be generated through increased translation initiation at downstream AUG sites, bypassing the CTSL signal peptide. Subsequent studies have shown that nCTSL regulates DNA repair [17], acts as a transcriptional regulator [18], and serves as a chromatin modifier [19]. In contrast, other reports have suggested that drug-induced nCTSL is a mediator of cancer cell death. Li et al. [20] demonstrated a correlation between 6-hydroxydopamine-induced nCTSL and aberrant expression of certain cell cycle regulators, leading to death of dopaminergic neurons. Dennemärker et al [21], utilizing mice with constitutive CTSL deficiency, reported that the absence of CTSL heightened tumor progression and metastasis. Moreover, knockdown of CTSL reduced rotenone-induced cell cycle arrest and cell death in PC-12 neuronal cells [22]. These pleotropic effects highlight the complexity of CTSL function in cancer biology, suggesting its effects are highly context-dependent and influenced by factors such as the type of cancer, the cellular microenvironment [23], and the presence of specific regulatory molecules or response to specific drugs. Importantly, previous studies have not examined a role for nCTSL in the action of CLF or examined possible synergy of CLF with PARP inhibitors in either hematological malignancies or solid tumors.

In the present study we observed that CLF induces a DNA damage response and enhances ovarian cancer sensitivity to PARP inhibitors, particularly olaparib. These effects, which are not seen with other nucleoside analogues such as cytarabine hydrochloride, fludarabine, or gemcitabine, are dependent on CLF induction of nCTSL. These observations identify a potentially promising combinatorial approach for extending PARP inhibitor action and an unexpected mediator of this effect.

## MATERIALS AND METHODS

### Materials

All anti-cancer drugs, inhibitors, reagents and antibodies are listed in Supplementary table 1.

### Cell culture and treatments

Details of cell lines and culture media used in the study are listed in Supplementary table 2. Cell lines were cultivated in respective culture media with 10% FBS, and 1% penicillin streptomycin in humidified incubators at 37 °C, 5% CO_2_ atmosphere. Current FDA-approved anticancer drugs were obtained from NCI-DCTD. The current set consists of 179 agents (10 mM) and details can be found here (https://dtp.cancer.gov/organization/dscb/obtaining/available_plates.htm). PEO1/PEO-OLTRC4 cells were treated at 10 µM concentrations for 24 hours and stained with Magic Red® Cathepsin L for observing CTSL activity. Further, based on our observation, nucleoside analogs including CLF, fludarabine, gemcitabine and cytarabine were used to treat PEO1 and PEO1-OLTRC4 cells for 24 hour. Cells were then fixed and stained with CTSL/CRM1 antibody to detect CTSL localization by confocal imaging. For clonogenic assays, 300 cells per well were seeded in 24-well plates and treated as indicated for up to 10 days till colonies became visible. The colonies were stained with crystal violet dye and quantified using the ImageJ software [24].

Clinical samples from chemo-naive patients were obtained at Mayo Clinic (IRB-approved protocol; 1288-03) and in association with the University of Minnesota Cancer Center Tissue Procurement Facility (IRB approval; 0702E01841). The patient ascites samples were centrifuged, and trypsinized for 10 min. Following further centrifugation, the cell pellet was treated for 5 min with 0.4% sterile ammonium chloride to eliminate the red blood cells and centrifuged. The pellets were then washed twice with sterile PBS and plated in DMEM/F12 medium supplemented with 15% FBS on low attachment plates. Briefly, 400 cells per well were plated in 96-well low attachment plates and treated with the indicated drug concentrations for up to 3 days. Cell viability was then measured using CellTiter-Glo® following the manufacturer protocol. Clinical characteristics of the patients, including the results of germline and/or somatic genetic testing that are part of standard care (e.g., FoundationOne CDx or Myriad myChoice CDx), are provided in the supplementary table 3.

### Subcellular fractionation studies

The cells treated with indicated doses of vehicle or CLF with/without PARP inhibitors *in vitro* and *ex vivo* were fractionated into the cytosol and nuclear portions using a NE-PER Kit (Thermo Fisher Scientific, USA) according to the manufacturer’s protocol. Briefly, the cells/tumor tissues were washed, incubated in cytoplasmic extraction solution and subsequently centrifuged. The supernatant served as the cytoplasmic fractions. The residual pellet was resuspended in nuclear extraction buffer, centrifuged for nuclear fractions separation.

### Generation of knockdown and overexpression cells

PEO1 and PEO1-OLTRC4 (Gift from Dr. Xinyan Wu, Mayo Clinic) cell lines were stably knocked down for CTSL using specific targeted shRNA from Sigma-Aldrich, following supplier’s protocol. CTSL-sh1 (GAATTGCCTCAGCTACTCTAA) CTSL-sh2 (TGCCTCAGCTACTCTAACATT) CTSL-sh3(AGGCGATGCACAACAGATTAT) and with nontargeted control shRNA as control with Lipofectamine 2000 (Invitrogen, Carlsbad, CA) as per manufacturer protocol in the indicated cell lines. Puromycin was used to select stable clones as reported earlier [25]. Snai1 shRNAs were kind gifts from Dr.Zhenkun Lou, Mayo Clinic, Rochester.

Flag-tag insert p3xFlag-CMV-14 was purchased from Sigma and human wildtype CTSL-C9 plasmid was purchased from Addgene (Addgene #11250). CTSL-wildtype construct was inserted into p3xFlag-CMV-14 to generate flag-tagged human hCTSL by using hCTSL-wildtype primers-CMV-14 Forward (Hind III) 5’ – AAT TAA GCT TAT GAA TCC TAC ACT CAT CCT TGC T-Reverse (Xba I)5’ – AAT TTC TAG ACA CAG TGG GGT AGC TGG CTG C-3’.

CTSL mutants were generated in human wildtype CTSL-C9 plasmid with the following replacements: AUG codons (Met 1, 35, 42 and 56) mutated to UUC (Phe) or UUG (Leu) to generate C9-tagged hCTSL-M1F, M35L, M42L and M56L by site-directed mutagenesis (Agilent) kit using primers hCTSL-M1F Fw-5’ C GGG CCC TCT AGA TTC AAT CCT ACA CTC ATC C 3’, R-5’ G GAT GAG TGT AGG ATT GAA TCT AGA GGG CCC G 3’; hCTSL M35L FW-5’ GG ACC AAG TGG AAG GCG TTG CAC AAC AGA TTA TAC GG 3’, RW-5’ CC GTA TAA TCT GTT GTG GAA CGC CTT CCA CTT GGT CC 3’; hCTSL M42L FW-5’C AAC AGA TTA TAC GGC TTG AAT GAA GAA GGA TGG AGG 3’ RW-5’ CCT CCA TCC TTC TTC ATT GAA GCC GTA TAA TCT GTT G 3’ hCTSL-M56L F-5’ GTG TGG GAG AAG AAC TTG AAG ATG ATT GAA CTG C 3’, R-5’ GCA GTT CAA TCA TCT TCA AGTTCT TCT CCC ACA C 5’; The M1F and M56L mutant constructs were overexpressed in the PEO1/OLTRC4 cells to generate nCTSL and stained with anti-C9 and confirmed by immunoblots. CTSL NLS mutant was generated by using WT-CTSL-Flag construct by site-directed mutagenesis (Agilent) kit using following primers F: 5’ CAG GTG ATG AAT GGC TTT CAA AAC GCT GCG CCC GCG GCG GGG AAA GTG TTC CAG GAA CCT 3’ and R:5’ AGG TTC CTG GAA CAC TTT CCC CGC CGC GGG CGC AGC GTT TTG AAA GCC ATT CAT CAC CTG 3’

### Cell Counting Kit-8 (CCK8) Assay

The Cells (PEO1/OLTRC4-shCTSL2 transfected with vector and M1F-CTSL) were seeded in 96-well plates and treated with CLF and olaparib at indicated concentration for 24 hours. Following treatment, 10 µL of CCK-8 reagent was added to each well and the cells were incubated for 4 h at 37°C and OD was estimated at 450 nm by a microplate reader. Cell proliferation was calculated relative to the untreated cells and analyzed using GraphPad Prism.

### Degradation of Cellular RAD51/53BP1 by rCTSL Analysis

To assess whether RAD51 and 53BP1 were directly degraded by CTSL, 50 µg of whole cell lysates from OVA-12 cells were incubated with recombinant human CTSL (rCTSL) protein (0.5 µg/sample for 1 hour) at room temperature in assay buffer (50 mM MES, 5 mM DTT, 1 mM EDTA, 0.005% (w/v) Brij35, pH 6.0). Samples were resolved on a 4-15% SDS-PAGE gel, followed by immunoblotting with the RAD51, 53BP1, PARP1 and tubulin antibodies [26]. Furthermore, human recombinant His-tagged RAD51 incubated with rCTSL or rCTSB for 1 hour in assay buffer in the presence of CTSL (z-fy(tbu)dmk) or CTSB (CA-074Me) specific inhibitors and followed by immunoblotting with the anti-His tag.

### Cellular thermal shift assay (CETSA)

PEO1 and PEO1/OLTRC4 cells were treated with indicated concentrations of DMSO, CLF and olaparib alone and whole cell lysates containing 0.1% Triton 100 were diluted to 1.5 μg/μl and the cell lysates were incubated for 30 min at room temperature. Proteins were denatured using Bio-Rad 96-well Thermal Cycler for 3 min at different temperatures ranging from 37 °C to 67 °C and centrifuged at 16000 X g for 30 min at 4 °C [27, 28]. The supernatants were immunoblotted and probed with anti-KPNB1 antibody and shown here.

### Single cell electrophoresis (COMET assay)

Alkaline comet assays were performed with the mentioned treatments using an established protocol [29]. 1×10^5^ cells were plated in 12 well plates and treated with CLF alone or in combination with olaparib or rucaparib for 24 hours. The cells were washed in 2 x PBS and resuspended in 100 µl of a low melting point agarose, smeared on glass slides. After gelling, slides immersed in pre-chilled alkaline lysis buffer (1.2 M NaCl, 100 mM Na2EDTA, 0.1% sodium lauryl sarcosinate, 0.26 M NaOH (pH > 13)) for 1 hour. Slides were electrophoresed in alkaline electrophoresis buffer (0.03 M NaOH, 2 mM Na2EDTA (pH ∼12.3)) for 15 min at 30 V. Slides were washed, stained with ethidium bromide (2.5 µg/ml) and observed using a fluorescence microscope (Olympus). Tail moments were quantified using CaspLab version 2.2 software and plotted.

### Homologous Recombination (HR) assay

PEO1 cells were stably transfected with the HR substrate pDR-GFP [30] and transfected with pCβASceI plasmid for 24 hours, and treated with indicated concentrations of CLF, olaparib alone and in combination for another 24 hours, and evaluated for the percent GFP positive fluorescence using flow cytometry.

### Immunoprecipitation assay

For detection of the protein complex, the cell lysates containing 400 μg of protein were incubated with the anti-Flag-CTSL antibody (1:100) overnight at 4 °C, and then 10 μl of 50% protein A-agarose beads were added and thoroughly mixed at 4 °C for 6 hours. The immunoprecipitates were washed thrice with PBS, and precipitated beads were loaded into the sample buffer, subjected to electrophoresis on 4– 15% SDS–PAGE and blotted using an anti-Flag,-KPNB1 or KPNA1 antibodies

### Immunofluorescence assay

Indicated cells were grown on multi-chambered slides overnight and then cells were fixed, permeabilized and stained with anti-RAD51,-γH2AX,-CRM1 and-CTSL (1:100) overnight at 4 °C. The cells were then stained using rabbit anti-mouse IgG Alexa Fluor® 594 or rabbit anti-mouse IgG-FITC (Molecular Probes) Slides were mounted with Antifade reagent (Invitrogen, Carlsbad, CA), visualized using a Zeiss-LSM780 confocal microscope and corrected total cell fluorescence (CTCF) was obtained by ImageJ-Fiji software.

### Immunoblot assay

Cytosolic and nuclear fractions or whole cell lysates were isolated and subjected to SDS-PAGE followed by immunoblot [31] using antibodies listed in Supplementary table 1. Secondary staining was achieved with secondary anti-mouse and rabbit with-680 or 800 IR dyes and scanned under the Odyssey Fc Imaging system (Bio-Rad, Hercules, CA). The normalized relative expression folds were calculated using ImageJ software.

### Annexin V/PI staining

Percent cell apoptosis was assessed as described before [26] and the cells after indicated treatments were fixed, stained with annexin V-pacific blue and propidium iodide (PI), then sorted using a flow cytometer (BD FACS Canto II) and analyzed using FlowJo 10.1 software

### Cell cycle analysis

Briefly cells were treated with the indicated concentrations of the mentioned drugs for 0-24 hours and the cells were collected, washed with ice-cold PBS, and fixed using 70% chilled alcohol and stained with propidium iodide in presence of RNase A (Invitrogen Life Technologies) at 37 °C for 20 min prior to analysis. Cell cycle was evaluated using FlowJo 10.1 software

### Synergy assessment

Drug combination effects were evaluated using Combenefit software (http://sourceforge.net/projects/combenefit/), which compares the observed drug response surface to a theoretical reference model and generates a percentage synergy score for each point in the dose-response matrix. Synergy and antagonism distributions were visualized, and drug combinations of CLF with rucaparib and olaparib (n = 3) were specifically assessed using the Loewe additivity model. Statistically significant interactions were highlighted. One sample t-test was used to assess the significance of observed synergy scores. In parallel, combination index (CI) values were determined using Calcusyn software (https://www.combosyn.com) based on the Chou–Talalay method, employing a non-constant ratio design [32].

### *Patient derived* xenograft (PDX) models

Female NSG mice (NOD-SCID) were injected intraperitoneally (i.p.) with ex-vivo cultures of PH610 and PH747 cells (4×10^6^ cells/mouse) and maintained for 15 days. The mice were then randomized into six groups (n=7, PH610) (n=6, PH747) and treated as; group 1: vehicle control; group 2: CLF (5 mg/kg every other day i.p.); group 3: olaparib (50 mg/kg/day by oral gavage); group 4: rucaparib (50 mg/kg/day by oral gavage); group 5: combination of CLF and olaparib; group 6: combination of CLF and rucaparib. Mice were treated for 4 weeks and then followed for 2 weeks and sacrificed when tumor burden exceeded 10% of body weight in the control group. At end of the study, tumor volume, ascites volume and survival of animals were assessed. Pairwise log-rank tests were performed for survival analysis and statistical significance (*p < 0.05; **p < 0.01; ***p <0.001). Before PDX model, body conditioning score was evaluated in SCID mice (n=4/group) by administering CLF (5mg/kg i.p 5d/w) alone in combination with rucaparib (50 mg/kg oral gavage daily) or olaparib (50mg/kg gavage daily) for 4 weeks. Animal experiments were carried out under the approved protocols and guidelines of the Mayo Clinic Animal Care and Use Committee (IACUC number#A00006931).

### KPCA (Syngeneic) tumor *model*

For animal studies, KPCA tumor model developed in C57BL/6 mice as described before [33]. In brief, three million cells were injected intraperitoneally (i.p.) and maintained for 7 days. The mice were then randomized into 4 groups (n=10), group 1: vehicle control; group 2: CLF (3.5 mg/kg 5d/w i.p.); group 3: olaparib (50 mg/kg/day by oral gavage); group 4: combination of CLF and olaparib; Mice were treated for 4 weeks and sacrificed when tumor burden exceeded 10% of initial body weight. At end of the study, tumor volume and survival of animals were assessed. Pairwise log-rank tests were performed for survival analysis and statistical significance (*p < 0.05; **p < 0.01; ***p <0.001). Animal experiments were carried out under the approved protocols and guidelines of the Mayo Clinic Animal Care and Use Committee (IACUC number# A00007937).

## Statistical analysis

All experiments were performed in triplicates in each of 3 independent experiments and the results were expressed as mean ± standard deviation unless otherwise noted. Statistical significance (*p < 0.05; **p < 0.01; ***p <0.001) was determined using Student’s test/GraphPad prism 10 unless otherwise noted.

## RESULTS

### CLF induces nCTSL in some OC cells

In the context of an ongoing effort to define the role of nCTSL, we conducted a drug repurposing screen with 179 FDA-approved drugs (https://dtp.cancer.gov/dtpstandard/ChemData/index.jsp) by immunofluorescence (IF) analysis of nCTSL in the *BRCA2*-mutant PEO1 ovarian cancer cell line and its isogenic olaparib resistant PEO1/OLTRC4 derivative. This screen showed that the anti-leukemic drug CLF promotes trafficking of CTSL to the nucleus in PEO1 cells, but not in PEO1/OLTRC4 cells (Figure 1A and B) respectively. For ease of presentation, we refer to cells in which CLF induces nCTSL as CLF-responsive (CLF-r) and those without nCTSL following CLF treatment as CLF-nonresponsive (CLF-nr). CLF promoted the nuclear accumulation of CTSL (Green foci in the nucleus) in PEO1 cells (CLF-r), both alone and in combination with olaparib or rucaparib (Figure 1C, top panel, CTSL: green, DAPI: blue). However, in PEO1/OLTRC4 cells (CLF-nr), only the CLF/olaparib combination (Figure 1D, bottom panel), but not CLF or rucaparib alone or in combination, promoted the nuclear accumulation of CTSL.

**Figure 1:**
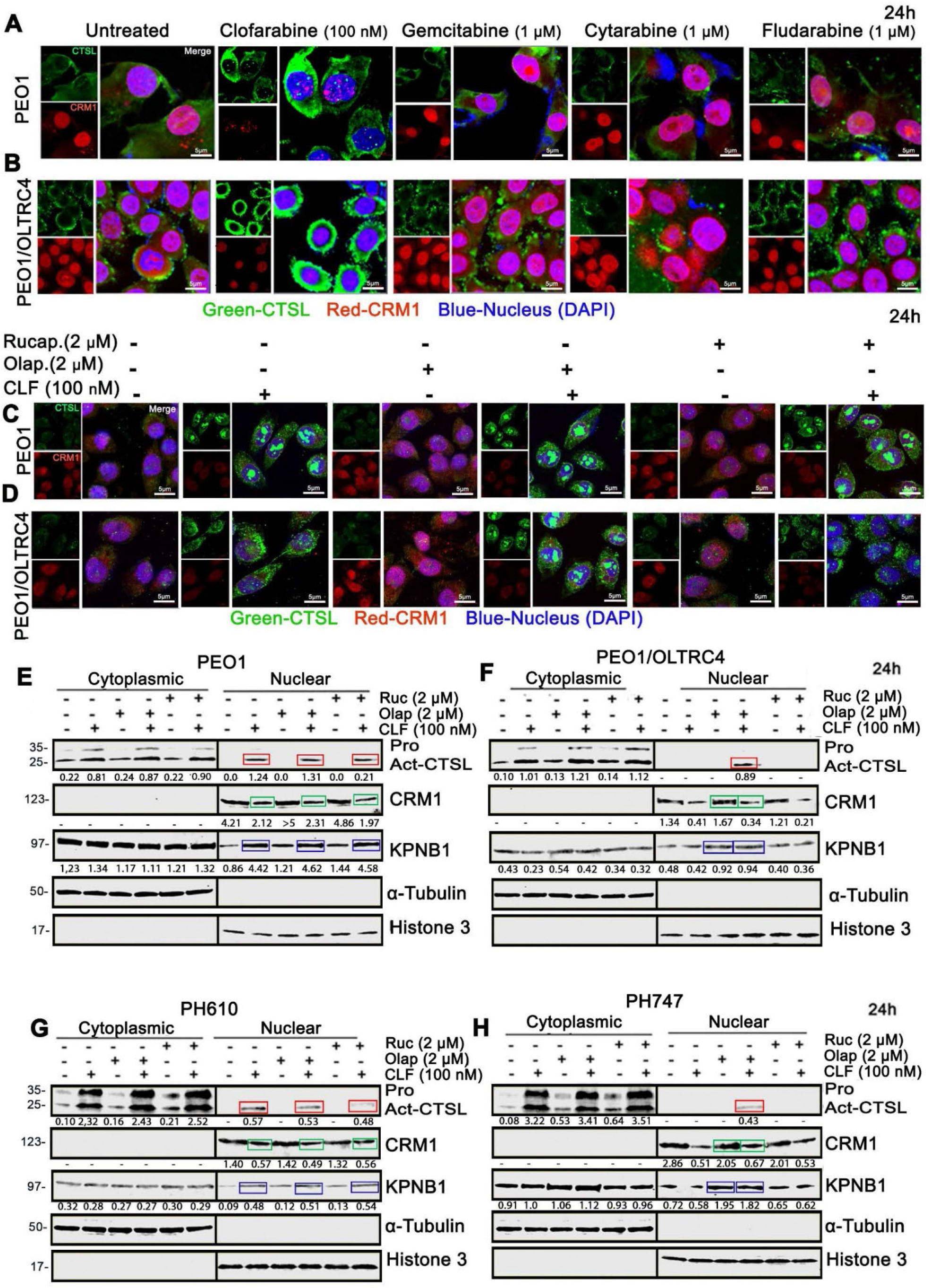
CLF induces differential nuclear trafficking of CTSL in CLF-r vs. CLF-nr ovarian cancer cells. (A) Representative immunofluorescence images of CTSL (green) and CRM1 (red) in PEO1 cells and (B) PEO1/OLTRC4 cells following 24-hours treatment with the indicated inhibitors are shown (40X magnification, scale bar: 5 μm). DAPI was used to stain the nucleus. (C) Similar immunofluorescence images were captured for PEO1 and (D) PEO1/OLTRC4 cells treated with CLF, either alone or in combination with the PARP1 inhibitors olaparib and rucaparib, at indicated concentrations for 24 hours (40X magnification, scale bar: 5 μm). Representative images include DAPI nuclear staining. (E) Western blot analysis of cytoplasmic and nuclear fractions from PEO1 cells and (F) PEO1/OLTRC4 cells were performed under similar treatment conditions. Tubulin was used as the cytosolic endogenous control, and Histone H3 served as the nuclear control. Normalized fold changes relative to untreated were calculated using ImageJ software are displayed beneath each panel. (G-H) Comparable immunoblot analysis of cytoplasmic and nuclear fractions was carried out on ex vivo patient-derived xenograft (PDX) cultures PH610 and PH747 under identical treatment conditions.

Western blotting indicated that CLF increases the amount of CTSL, particularly a lower molecular weight isoform and aids in the nuclear accumulation of CTSL in the PEO1 cells, alone and in combination with olaparib or rucaparib (Figure 1E, red box). In contrast, in the PEO1/OLTRC4 cells, only the combination of CLF and olaparib, but not rucaparib, resulted in the accumulation of this CTSL isoform in the nucleus (Figure 1F, red box). Additionally, CLF induced nCTSL in *ex vivo* culture of a patient-derived xenograft PH610 cells (BRCA WT, TP53 mutation not tested-Supplementary table 3) (Figure 1G, red boxes) classifying them as CLF-r cells, but was not sufficient to induce nCTSL in PH747 (BRCA1 and TP53 mutant-Supplementary table 3) cells (Figure 1H, red box) which required a combination of CLF+olaparib to aid in nCTSL translocation, classifying them as CLF-nr cells. Similarly, CLF alone aided in nCTSL accumulation in OVCAR5 and HeyA8 cells (all BRCA WT), categorizing them as CLF-r. In contrast, CLF alone was not sufficient to induce nCTSL in OVCAR8 and HeyA8-MDR (with cross-resistance to cisplatin, adriamycin, and vincristine) [34], thus classifying them as CLF-nr. These data suggest that the restriction on trafficking of nCTSL with CLF monotherapy is not limited to PARP inhibitor-resistant cells but also extends to cells resistant to paclitaxel, cisplatin, and other drugs (Supplementary table 4).

In further experiments, the PARP inhibitors niraparib and veliparib, like rucaparib, failed to induce nCTSL alone or in combination with CLF in CLF-nr PH747 cells (Supplementary figure 1A-C). Additional studies confirmed differences in CLF-induced accumulation of nCTSL between samples. In particular, CLF induced nCTSL in *ex vivo* cultures of patient-derived ascites OVA2 but not in OVA12 (Supplementary figure 1D and E), and in *ex vivo* cultures of patient primary tumor OV-1182 but not in OV-1183 (Supplementary figure 1F and G).

### CLF and olaparib differentially regulate the trafficking of Importin B1 (KPNB1) to the nucleus in the CLF-r vs. CLF-nr OC cells

Considering that nuclear proteins are imported by KPNB1 and exported by Exportin 1 (also known as CRM1), we assessed the extent to which CLF and PARP inhibitors affected the expression of these proteins. In CLF-r PEO1 cells, CLF downregulates CRM1 and upregulates KPNB1 (Figure 1E). In CLF-nr PEO1/OLTRC4 cells, CLF downregulates CRM1 but fails to upregulate KPNB1 (Figure 1F). These results, which are consistent with prior studies indicating that KPNB1 knockdown (KD) resulted in the absence of nCTSL in the nucleus of MDA-468 cells [35], suggest that KPNB1 upregulation determines nuclear CTSL import.

In PEO1 and PH610 cells, CLF (but not olaparib) promoted the trafficking of KPNB1 into the nucleus (Figure 1E and G, blue boxes), as seen as increased levels of KPNB1 in the nuclear compartment. In contrast, in the CLF-nr PEO1/OLTRC4 and PDX PH747 cells, olaparib (but not rucaparib nor CLF) regulated KPNB1 trafficking to the nucleus (Figure 1F and H, blue boxes vs. last two lanes). CLF downregulated olaparib-induced CRM1 (Figure 1F and H, green boxes), leading to the retention of CTSL in the nuclear compartment. Furthermore, it is important to note that rucaparib (as well as niraparib and veliparib, data not shown) alone or in combination with CLF do not promote KPNB1 nuclear import in CLF-nr cells (Figure 1F and H, last two lanes).

### Importazole specifically blocks CLF-or CLF+olaparib-induced KPNB1 nuclear import

Our data suggests the possibility that, in CLF-r cells, CLF facilitates the nuclear translocation of KPNB1-bound CTSL, while in CLF-nr cells, olaparib aids this process. Further experiments were performed to test this model. The first of these experiments were designed to assess whether KPNB1 and CTSL interact. When Flag-tagged full length CTSL was expressed in HEK293 cells, coimmunoprecipitation using anti-KPNB1 or anti-Flag antibodies showed that KPNB1, but not KPNA1, pulled down with CTSL and vice versa (Figure 2A).

**Figure 2:**
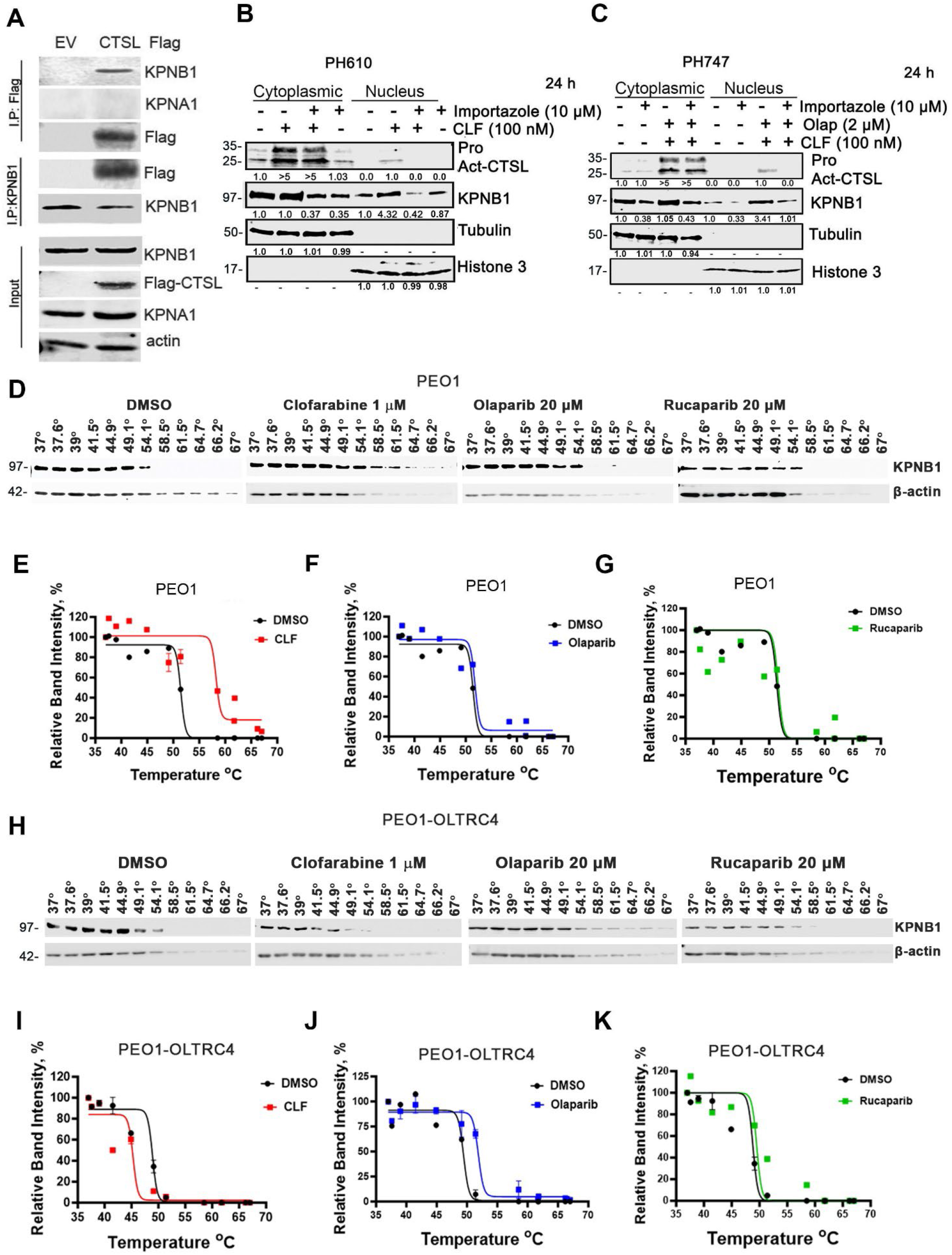
Importazole specifically blocks CLF alone and CLF+olaparib-induced KPNB1 import into the nucleus. (A) Cell extracts from empty vector and Flag-tagged CTSL overexpressing cells were immunoprecipitated with an anti-Flag antibody. Co-precipitated KPNB1 was detected via immunoblot analysis, and the reverse co-immunoprecipitation was also performed. KPNA1 was included as negative control. (B-C) Western blot analysis of CTSL and KPNB1 levels was performed on cytoplasmic and nuclear fractionated extracts from PH610 and PH747 cultures treated with the indicated drugs with or without importazole for 24 hours. Normalized fold changes relative to untreated were calculated using ImageJ software. (D) CETSA assays were conducted to evaluate the thermal stability of KPNB1 protein upon binding to CLF or olaparib/rucaparib in PEO1, followed by analysis via western blot. (E-G) Densitometric analysis of the CETSA blot from PEO1 cells was performed using ImageJ. (H-K) Similar type of CETSA experiments conducted in PEO/OLTRC4 and relative CETSA band intensity was analyzed by ImageJ.

Next, to determine whether KPNB1 is involved in the nuclear translocation of CTSL, we treated CLF-r PH610 and CLF-nr PH747 cells with CLF or olaparib, alone or in combination (Figure 2B and C), and examined fractionated cell lysates by western blot. In PH610 cells, CLF alone induced increased levels of KPNB1 in the nuclear compartment as well as nuclear CTSL (nCTSL) (Figure 2B). In cells treated with importazole, a small molecule inhibitor that specifically blocks KPNB1-mediated nuclear import without disrupting CRM1-mediated nuclear export [36], the levels of nuclear KPNB1 were negligible (Figure 2B), coinciding with the absence of nCTSL (Figure 2B). Similar importazole-inhibitable translocation of KPNB1 as well as CTSL was observed in CLF-nr PH747 cells treated with the CLF + olaparib combination (Figure 2C). Collectively, these results support the role of KPNB1 as the importer involved in the translocation of nCTSL.

### Cellular thermal shift assay shows olaparib stabilizes KPNB1 bound CTSL

To better understand the differential role of CLF versus olaparib in import of CTSL-bound KPNB1, we performed a cellular thermal shift assay (CETSA) [27, 28] to assess the impact of these drugs on KPNB1 thermal stability. In CLF-r PEO1 cells, KPNB1 is stabilized by CLF, as evidenced by its higher denaturation temperature compared to the untreated cells, (Figure 2D, second vs. third or fourth panel). There is a notable increase in the stability of KPNB1 bound to CLF but not to olaparib or rucaparib in these cells, as shown by the melting curves with relative band intensities in Figures 2E - G (compare the red and blue or green lines). Conversely, in PEO1/OLTRC4 cells, KPNB1 is stabilized by olaparib but not by CLF or rucaparib (Figure 2H, second vs. third or fourth panel). A thermal shift is observed with olaparib but not CLF in these cells, as depicted by the melting curves in Figures 2I-K (compare the red and blue lines). No thermal shift was seen with rucaparib in both CLF-r and CLF-nr cell lines. The control actin profile is similar between the two drugs and cell lines. These data support the differential stability of CTSL-bound KPNB1 translocated to the nucleus by clofarabine in CLF-r PEO1 cells and by olaparib in CLF-nr PEO1/OLTRC4 cells, respectively.

### Drug-induced CTSL nuclear translocation occurs in cancer cell lines of different origins

To assess whether CLF-or CLF + olaparib-induced nuclear translation of CTSL occurs in other cell types, we explored a number of additional cancer cell lines (Supplementary table 5, Supplementary figure 2A-E). In the four triple negative breast cancer lines examined (MDA-MB-231, HCC1395, MDA-MB-436, and HCC1937), CLF alone facilitated nuclear CTSL trafficking, classifying them as CLF-r cells. Conversely, in the luminal breast cancer cell line MCF7 cells, CLF + olaparib promoted CTSL nuclear trafficking, categorizing them as CLF-nr cells. Similarly, the OV2008 cervical cell line was CLF-r, whereas isogenic cisplatin-resistant C13 cells were CLF-nr, mirroring the behavior of ovarian cell lines HeyA8 and HeyA8 MDR cells. The endometrial cancer cell lines KLE and ARK1 were CLF-nr. The uterine carcinosarcoma cell line SNU539 was CLF-r, while SNU1077 was categorized as CLF-nr. Non-small cell lung cancer cell lines H460 and A549 also belonged to the CLF-nr category. Collectively, the data suggests that CLF is able to promote nuclear CTSL translocation in a subset of cell lines of different histological origins, whereas the CLF+ olaparib combination is able to promote nuclear CTSL translocation quite broadly.

### Nuclear CTSL targets 53BP1 and RAD51 to inhibit homologous recombination repair

Based on reports that nCTSL degrades 53BP1 in breast cancer [17], we evaluated the ability of drug-induced nCTSL to target two major repair proteins, 53BP1 and RAD51. In PEO1 and PEO1/OLTRC4 cells (and in PH610 and PH747, data not shown), the presence of drug-induced nCTSL is associated with reduced levels of 53BP1 and RAD51 in both CLF-r and CLF-nr cells (Figure 3A and B, respectively). Consistent with these findings, IF analysis revealed that CLF downregulated 53BP1 and RAD51 in PEO1 non-targeted control (NTC) cells, but not in CTSL-knockdown (KD) sh2 and sh3 cells (Figure 3C), indicating that nCTSL plays a role in degrading these specific proteins. Figures 3D-F show the quantification of the number of 53BP1 and RAD51 foci in PEO1 NTC and CTSL-KD sh2 and sh3 cells, respectively. Data using recombinant CTSL (rCTSL) incubated with lysates from OVA-12 cells confirmed that, in addition to 53BP1, CTSL also reduced RAD51 (Figure 3G) but not PARP1. Treatment with the CTSL-specific inhibitor z-fy(tbu)-dmk restored 53BP1 and RAD51 levels without affecting PARP1, suggesting that CTSL selectively degrades certain DNA repair proteins.

**Figure 3:**
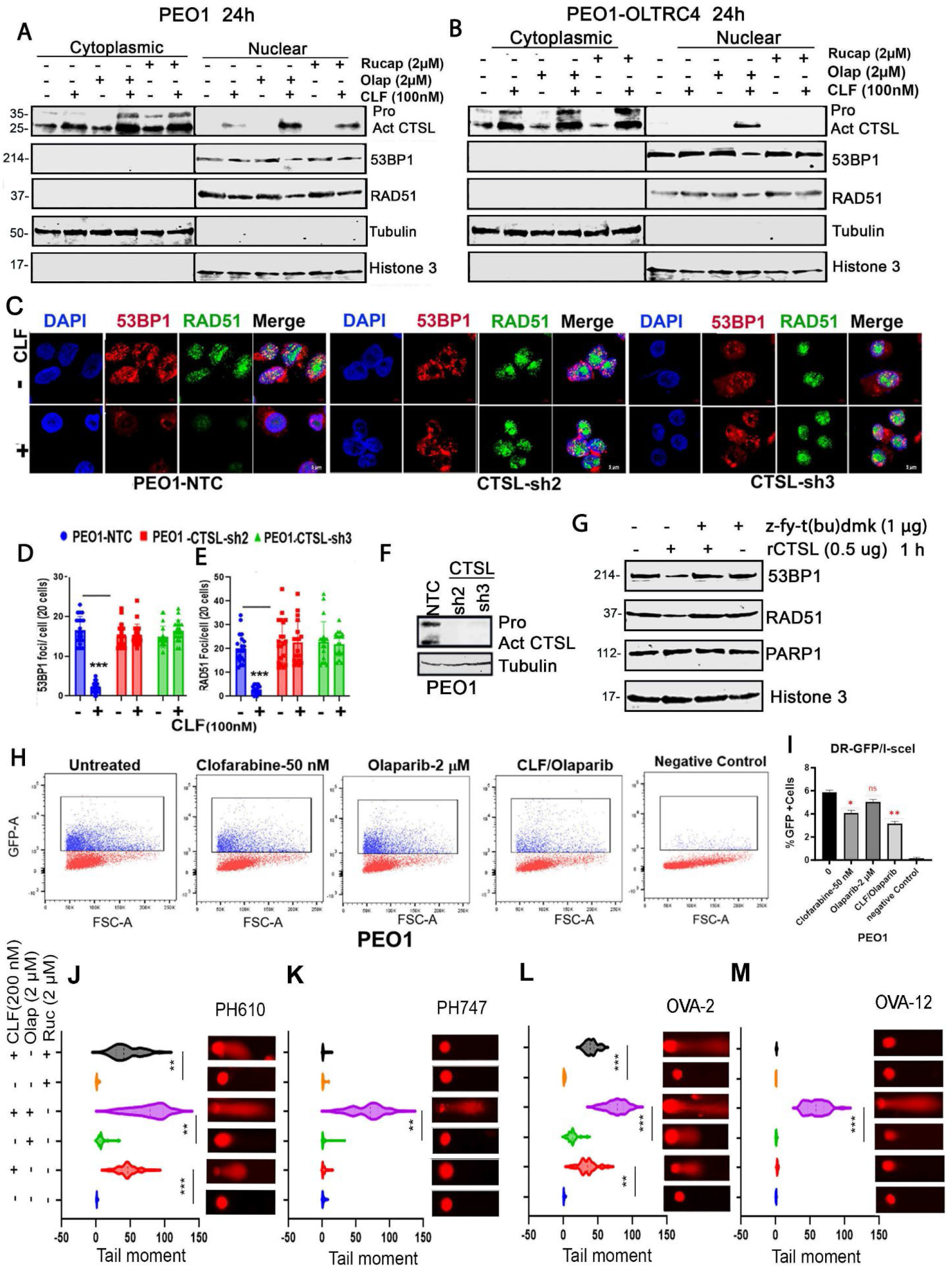
nCTSL targets 53BP1 and RAD51 to inhibit the homologous recombination repair and induce DNA damage. (A-B) Immunoblot analysis was performed to evaluate CTSL, 53BP1, and RAD51 levels in cytoplasmic and nuclear fractions of PEO1 and PEO1/OLTRC4 cells treated with the indicated drugs for 24 hours. Tubulin was used as the cytosolic control, and Histone H3 as the nuclear control. (C) Representative immunofluorescence images show 53BP1 (red) and RAD51 (green) foci in PEO1 non-targeting control (NTC), CTSL KD sh2, and sh3 cells following 24-hour CLF treatment. DAPI was used to stain nuclei. (D-E) Quantification of the number of foci per cell was performed in 25 cells, and data were plotted as mean ± SEM (***p < 0.001 vs. control). (F) Immunoblot analysis confirmed efficient knockdown (KD) of CTSL in PEO1 cells. (G) Immunoblot analysis of RAD51, 53BP1, and PARP1 levels was conducted on OVA12 ascites cell lysates treated with recombinant CTSL protein alone or in combination with the z-t(bu)-dmk inhibitor for 24 hours. (H) Representative flow cytometric analysis images show the percentage of GFP-positive cells as a measure of double-strand break (DSB) repair following treatment with the indicated concentrations of CLF and olaparib, either alone or in combination, in PEO1 cells for 24 hours. (I) Quantification of GFP-positive cells is presented as mean ± SD (*p < 0.05, **p < 0.01, ***p < 0.001 vs. control). (J-K) Representative images from the alkaline COMET assay are shown for PDX PH610 (CLF-r) and PH747 (CLF-nr) cells treated with CLF (200 nM) and olaparib (2 µM) or rucaparib (2 µM), either alone or in combination, for 24 hours. Tail moments of about 50 comets from each untreated and treated groups were measured by casplab software and graphed (*p < 0.05, **p < 0.01, ***p < 0.001 vs. control or monotherapy as indicated). (L-M)) A similar COMET assay was performed in ovarian ascites cell lines OVA-2 and OVA-12 treated with olaparib (2 µM), and rucaparib (2 µM), either alone or in combination with CLF, for 24 hours (*p < 0.05, **p < 0.01, ***p < 0.001 vs. control or monotherapy).

Thus, we have identified RAD51 as an additional DNA repair protein targeted by CTSL. To validate this, His-tagged recombinant RAD51 (rRAD51) was incubated with recombinant CTSL (rCTSL) or CTSB (rCTSB) for 1 hour. Western blot analysis showed degradation of His-tagged rRAD51 by rCTSL and this degradation was inhibited by the CTSL-specific inhibitor z-fy(tbu)-dmk, while CTSB had no effect under the same conditions (Supplementary figure 3A-B).

Additionally, PEO1/ OLTRC4 cells treated with the CLF/olaparib combination in the presence of either the proteasome inhibitor MG132 or the CTSL-specific inhibitor z-fy(tbu)-dmk showed that nCTSL is responsible for the degradation of RAD51 and 53BP1. Notably, this degradation was blocked only by z-fy(tbu)-dmk, but not by MG132, indicating that CTSL mediates this effect independently of the proteasome. Inhibition of CTSL with z-fy(tbu)-dmk restored RAD51 and 53BP1 levels following CLF/olaparib treatment, further supporting a direct role for nCTSL in regulating RAD51 stability (Supplementary figure 3C).

To assess the effects of CLF and olaparib, individually or in combination, on homologous recombination repair (HRR), we employed the DR-GFP reporter assay [37]. In PEO1 (CLF-r) cells expressing the DR-GFP substrate, I-SceI-induced double-strand breaks resulted in a significant reduction in GFP⁺ cells following CLF or CLF+olaparib treatment-both of which promote CTSL nuclear translocation, compared to olaparib alone or untreated controls, indicating impaired HRR (Figures 3H-I). Contrary to previous reports suggesting that nuclear CTSL (nCTSL) promotes HRR by degrading 53BP1, our data show that nCTSL concurrently targets both 53BP1 and RAD51, thereby blocking the restoration of HR repair. These findings suggest that the CLF+olaparib combination impairs DNA repair by downregulating two critical HRR factors, 53BP1 and RAD51.

### Prescence of nuclear CTSL is associated with DNA damage in PDX and OVA cells

Concomitant with the induction of nCTSL in CLF-r PH610 and OVA-2 cells, COMET assays demonstrated that CLF alone or in combination with olaparib induced substantial DNA damage, whereas olaparib alone induced negligible DNA damage (Figures 3J and L). In contrast, CLF alone produced negligible DNA damage in CLF-nr PH747 and OVA-12 cells, but the combination of CLF and olaparib (but not rucaparib) induced substantial DNA damage in these cells (Figures 3K and M). Quantification of tail moments (>30 comets/group) for all events confirmed the visual impression of the COMET images. Importantly, CTSL knockdown markedly inhibited the ability of CLF alone or in combination with olaparib to induce DNA damage compared to PEO1 cells transduced with non-targeting control (NTC) (Supplementary figure 4A-B). Collectively, these results show that DNA damage induced by these drugs rely on the presence of CTSL in the nucleus.

### CLF+olaparib combination induces G2M arrest and apoptosis in CLF-nr cells

To better understand the effect of DNA damage induced by CLF alone or in combination with olaparib or rucaparib, CLF-r (PEO1 & PH610) and CLF-nr (OLTRC4 & PH747) cells were treated with 200 nM CLF alone or in combination with 2.0 µM olaparib or rucaparib for 24 hours. Cell cycle analysis revealed that CLF alone or in combination with PARP inhibitors led to G2/M phase cell cycle arrest of CLF-r PEO1 cells (Figure 4A), whereas only the combination of CLF and olaparib resulted in G2/M arrest CLF-nr PEO1/OLTRC4 cells (Figure 4B) and furthermore, similar type of G2/M arrests were observed in *ex vivo* cultures of PH610 and PH747 at the same treatment conditions (Supplementary figure 4C-F). Consistent with the cell cycle analysis, the apoptotic effects of CLF were examined in both the CLF-r PEO1 and CLF-nr PEO1/OLTRC4 models using flow cytometry with Annexin V (Pacific Blue) and propidium iodide labeling with and without the caspase-3 specific inhibitor Z-DEVD-FMK. In PEO1 cells, CLF alone induced ∼20% cell death, and this was enhanced more than 4-fold when combined with olaparib, with minimal attenuation by the caspase-3 inhibitor (Figure 4C and D). These data suggest that the caspase-independent cell death as a mechanism in these cells. Similarly, in CLF-nr cells, CLF+ olaparib induced cell death of >80% of cells with minimal attenuation in the presence of Z-DEVD-FMK, again implicating a caspase-independent mechanism in nCTSL-mediated cell death (Figure 4E and F). Since CLF induces caspase-independent cell death, we first validated the efficacy of the caspase-3 inhibitor Z-DEVD-FMK. PEO1 cells were treated with doxorubicin-a known inducer of caspase-dependent apoptosis [38], alone or in combination with Z-DEVD-FMK. Doxorubicin treatment led to the cleavage of caspase-3 and PARP1, whereas co-treatment with Z-DEVD-FMK effectively blocked caspase-3 activation and PARP1 cleavage, confirming the functional efficacy of the Z-DEVD-FMK. (Supplementary figure 4G). Collectively, these data indicate that CLF and olaparib combination promotes nCTSL translocation and induces caspase-independent cell death in both CLF-r and CLF-nr cells. The observed cell death is predominantly caspase-independent, consistent with replication stress–induced catastrophic DNA damage rather than classical apoptosis. Combined with the protective effect of CTSL knockdown (Supplementary figure 4A-B), these observations provide evidence for the role of nCTSL in the DNA damage and cell death.

**Figure 4:**
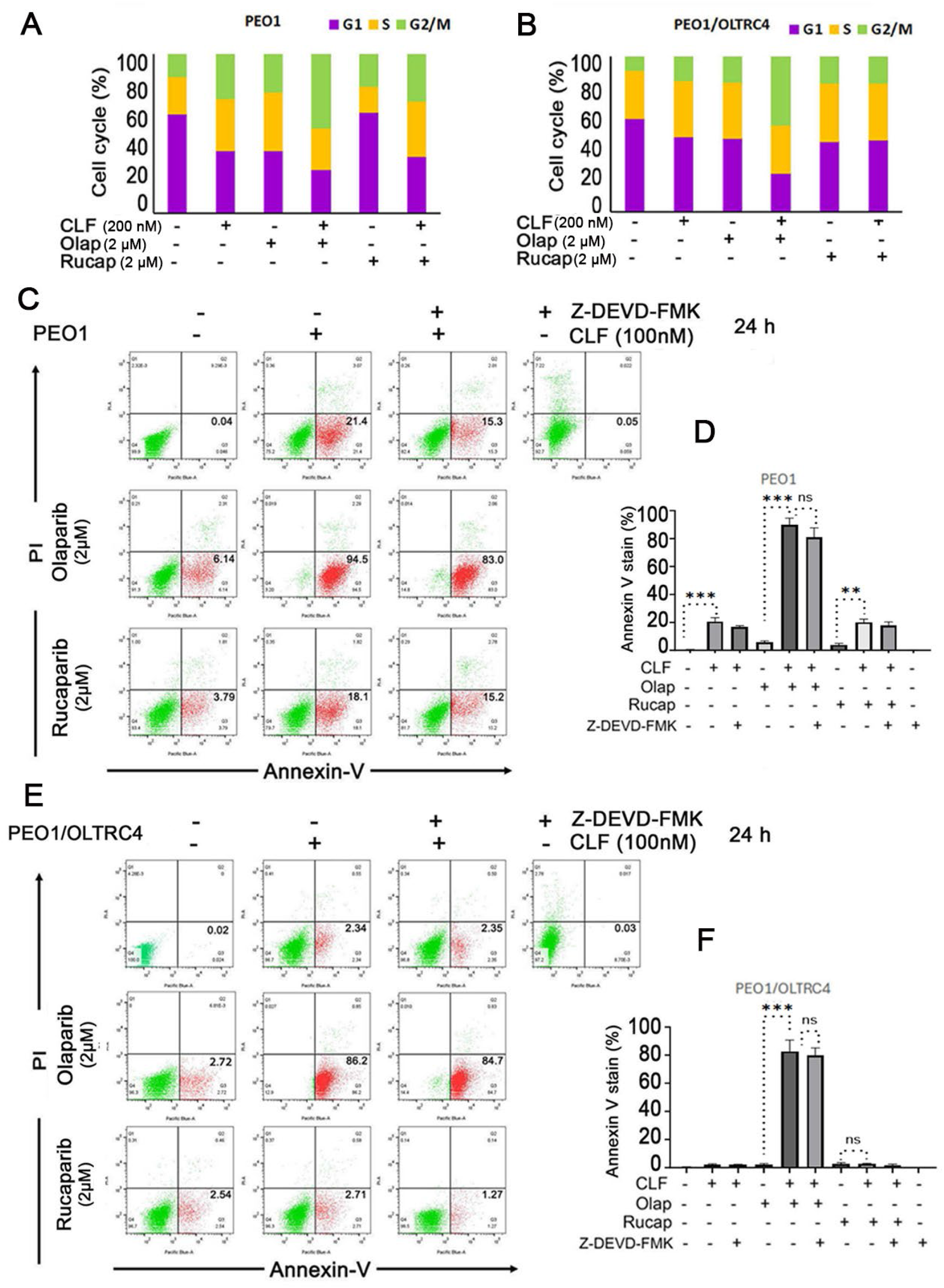
Combination of CLF with olaparib treatment induces G2M arrest and apoptosis in CLF-nr cells. (A) Cell cycle analysis of PEO1 cells was performed by PI staining and flow cytometry following ex vivo treatment with the indicated concentrations of CLF, olaparib, or rucaparib, either alone or in combination, for 24 hours. The percentage of cells in each cell cycle phase was quantified. (B) Similar cell cycle analysis was conducted in PEO1/OLTRC4 cells. (C) Annexin V/PI stain and flow cytometric analysis of apoptosis in PEO1 cells treated with CLF, alone or in combination with olaparib or rucaparib, in the presence or absence of the caspase 3 inhibitor Z-DEVD-FMK (5 µM) for 24 hours. (D) Quantification of Annexin V– positive PEO1 cells. (E) Similar cell cycle analysis was conducted in PEO1/OLTRC4 cells. (F) Quantification of Annexin V–positive PEO1/OLTRC4 cells. Data are presented as mean ± SD (*p < 0.05, **p < 0.01, ***p < 0.001 vs. control).

### CLF synergizes with olaparib and rucaparib in CLF-r cells and only with olaparib in CLF-nr cells

Given the differential ability of CLF to induce nCTSL in CLF-responsive versus CLF-nonresponsive cells, we next examined whether the presence of nCTSL is required for CLF-mediated sensitization to PARP inhibitors. Clonogenic assays of PEO1 and its isogenic olaparib-resistant PEO1/OLTRC4 cells showed that PEO1 cells were more sensitive to olaparib compared to PEO1/OLTRC4 cells (Figure 5A and B). The IC50s for olaparib and rucaparib in CLF-r PEO1 cells decreased 10-fold (from 1.8 µM to 0.18 µM) and 4-fold (from 0.4 µM to 0.1 µM), respectively, when 5.0 nM CLF was included (Figures 5C and D). The changes were smaller in CLF-nr PEO1/OLTRC4 cells, with a 3-fold decrease in olaparib IC50 from 11.76 µM to 3.5 µM when 5.0 nM CLF was included (Figure 5E) and smaller decrement in rucaparib IC50 in combination with CLF (Figure 5F, blue line).

**Figure 5:**
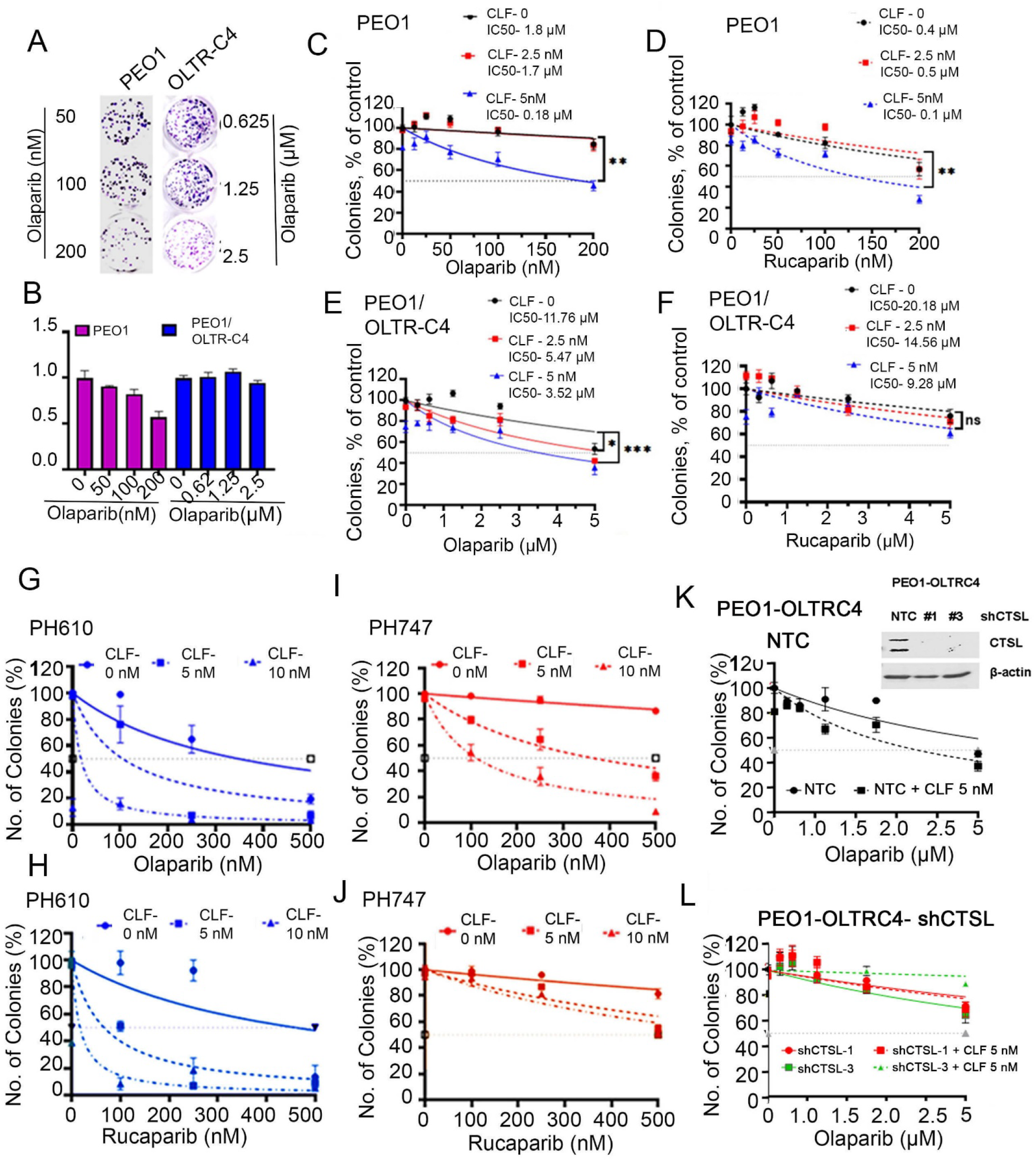
CLF-r cells show synergy on combination treatment of CLF with olaparib and rucaparib, while CLF-nr cells show similar effect with CLF + olaparib only. (A) Representative images of the colony formation assay are shown in PEO1 and PEO1/OLTRC4 cells with increasing dose of olaparib as indicated. (B) Quantification was performed using image J software and plotted. (C-D) Quantification of the clonogenic assay was represented as colonies (% of the control) in the PEO1 cells upon continuous exposure to increasing concentrations of olaparib and rucaparib respectively (±) 2.5 and 5.0 nM CLF. (E-F) Similar assay was performed and plotted in the PEO1/OLTRC4 cells. (G-H) Similar assay was performed with the indicated drug concentrations in the PDX PH610 and (I-J) PH747 cells respectively. (K-L) Quantification of the clonogenic assay was represented as colonies (% of the control) in the PEO1/OLTRC4 NTC and CTSL-KD cells upon continuous exposure to increasing concentrations of olaparib with 5 nM of CLF (Inset, western blot shows shCTSL1 and shCTSL3 knockdown compared to NTC). For all the cases, IC50 value was calculated and presented (error bars, ±SEM, n=3).

Consistent with the cell line model data, clonogenic assays of cultures ovarian cancer PDXs grown *ex vivo* showed that CLF sensitized the CLF-r PDX PH610 to both olaparib and rucaparib (Figure 5G and H), whereas CLF sensitized the CLF-nr PDX PH747 cells to olaparib but not to rucaparib (Figure 5I and J). Once again nCTSL was necessary for sensitization, as CTSL knockdown rendered PEO1/OLTRC4 cells resistant to olaparib alone or combination with CLF compared to controls transduced with nontargeting shRNA (Figure 5K and L).

### Met56 is essential for nuclear localization of CTSL

The 5′ portion of the human cathepsin L cDNA contains 7 AUG codons, corresponding to methionine 1, 35, 42 56, 58, 75, 77, 83, and 92. hCTSL mutants were generated with the following substitutions in AUG codons (Met 1, 35, 42 and 56). We have generated several C9-tagged CTSL constructs including with substitutions in the first initiation codon ATG > TTC, converting Methionine to Phenylalanine-referred to as M1F mutant and ATG > TTG, converting Methionine to Leucine-referred to as M35L, M42L, M56L mutant (Supplementary figure 5A) [39]. M1F-Mut prevents the generation of the secreted form of CTSL [16]. To validate the specific role of nCTSL in DDR, we transfected WT, M1F and M56L [16] into PEO1/OLTRC4 sh2CTSL cells, which have low levels of endogenous CTSL due to shRNA-mediated silencing. IF analysis detected nCTSL-C9 in cells transfected with M1F but not with the M56L (Supplementary figure 5B), a finding corroborated by cell fractionation and immunoblot analysis as well (Supplementary figure 5C). Expression of the M35L and M42L mutant constructs, the two other in-frame start codons in the human CTSL in the M1F-Mut background, localized to the nucleus implicating codon 56 (M56) as the preferred first in frame ATG to generate the nCTSL isoform. The presence of nCTSL in the transfected cells was again accompanied by enhanced DNA damage as assessed by increased γH2AX levels as well as downregulated 53BP1 and RAD51 (Supplementary figure 5C). Furthermore, to establish the fundamental role of nCTSL, we performed a genetic rescue experiment in OLTRC4 CTSL KD cells. Ectopic M1F-CTSL expression restored sensitivity to CLF and olaparib and enhanced the efficacy and synergistic interaction of the CLF/olaparib combination compared with vector control (Supplementary figure 5D-F). These findings provide genetic evidence supporting the role of intracellular/nuclear CTSL in CLF-mediated PARP inhibitor sensitization.

Previous studies reported that Snail promotes nuclear localization of CTSL and Cux1 through its nuclear localization signal (NLS) via KPNB1 [16, 40]. To determine whether Snail contributes to nuclear CTSL (nCTSL) accumulation in our system, Snail (Snai1) was silenced using shRNAs, and efficient knockdown was confirmed by Western blotting (Supplementary figure 6A). CLF and olaparib treatment markedly reduced Snail expression in both cytosolic and nuclear fractions of NTC-OLTRC4 cells, with further reduction observed in Snail-knockdown cells. However, Snail depletion did not alter CLF/olaparib-induced nuclear trafficking of CTSL or KPNB1 (Supplementary figure. 6B). Consistent with this observation, confocal microscopy showed that NLS-mutant CTSL failed to translocate to the nucleus, whereas wild-type CTSL containing an intact NLS efficiently localized to the nucleus (Supplementary figure. 6C–D), indicating that CTSL relies on its intrinsic NLS for nuclear import. Functionally, cells expressing NLS-mutant CTSL did not undergo cell death, while cells expressing wild-type NLS-CTSL exhibited significant cell death, as determined by Annexin V/PI staining (Supplementary figure. 6E-F). Together, these findings demonstrate that CLF and olaparib promote nuclear accumulation of CTSL independently of Snail, and that CTSL nuclear trafficking and associated cell death require its intrinsic NLS.

### Evaluation of synergy between CLF and olaparib or rucaparib in clinical samples ex vivo

To assess the potential efficacy of the CLF and PARPi combination in primary OC samples, we examined 14 patient-derived ascites samples (Supplementary table 3). OC cells were enriched in these cultures based on epithelial marker PAX8 expression, with almost no fibroblast activated protein (FAP) expression (Supplementary figure 7A). To evaluate the effect of CLF alone or in combination with olaparib or rucaparib, spheroids were generated (Supplementary figure 7B) with OVA-11 and OVA-12 ascites using 12.5% ascitic supernatant from the corresponding patients. Supplementary figure 7C shows the efficacy of the drugs expressed as spheroid area compared to untreated control.

The calculated Combination Index (CI) values for the individual drug combinations across three categories (CLF-r, CLF-nr and CLF-res as defined in this study) of 14 ascites samples are shown in Figure 6 and supplementary fig. 8. Six CLF-r ascites samples (OVA-1,-2,-10,-11,-22, and-33) (Figure 6A) and four CLF-nr ascites samples (OVA-12,-15,-28 and-34) (Figure 6C), collectively representing ∼72% (10/14) of the ascites, displayed synergism with the CLF+olaparib combination, with CI values ranging between 0.4 and 0.7 (Figures 6D). In the CLF-r OVA samples, CLF also synergized with rucaparib, with CI values ranging from 0.38 to 0.7 (Figure 6B). In contrast, in the CLF-nr OVAs, the CLF+rucaparib combination was antagonistic (Figure 6D). In the other 28% (4/14) of the ascites samples (OVA-9,-16, - 29, and-30), the CLF+olaparib combination was antagonistic, categorizing them as CLF-Res (resistant) cells (Supplementary figure 8A-B).

**Figure 6:**
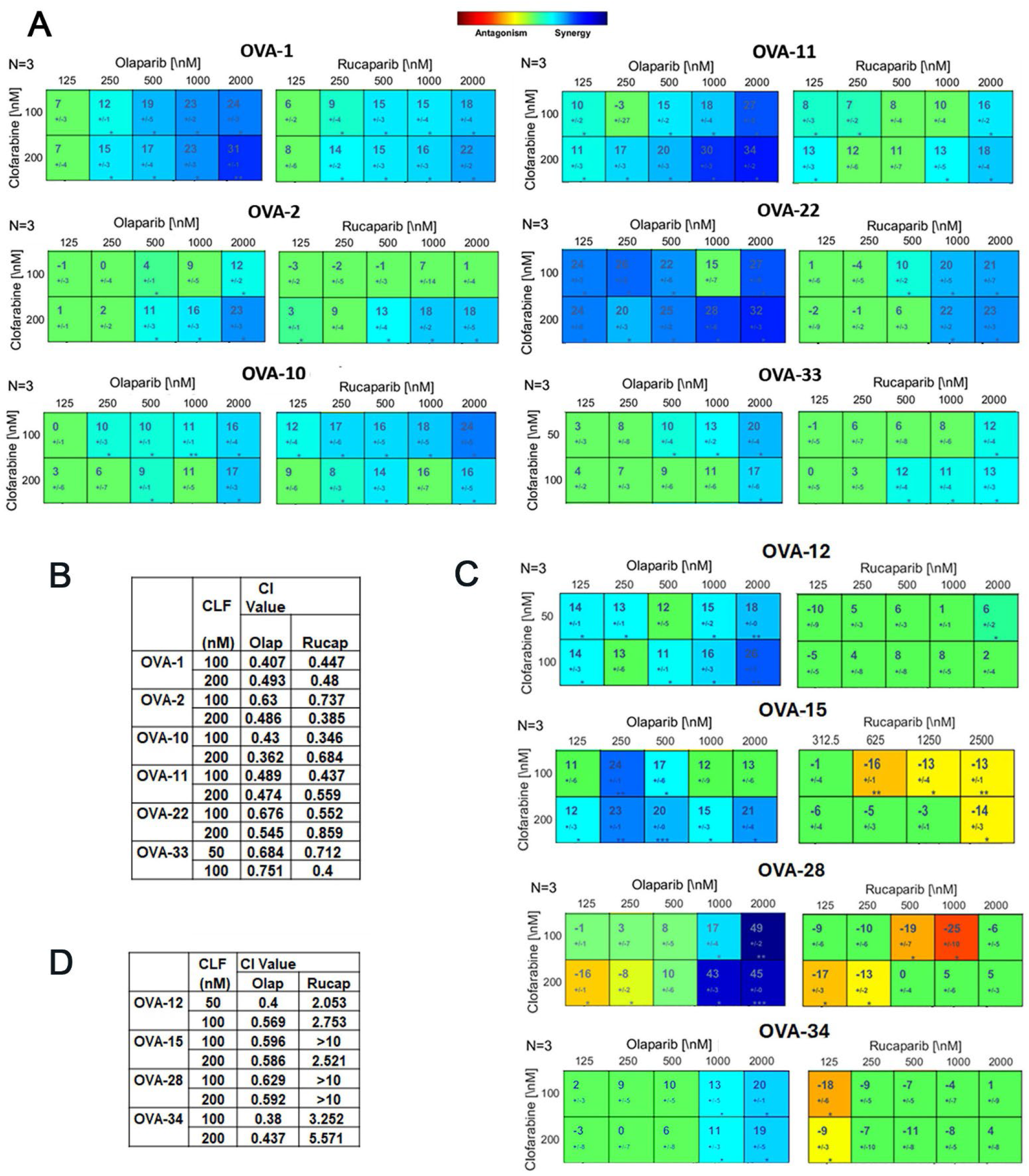
Evaluation of synergy upon treatment of CLF with olaparib and rucaparib respectively in the ascites-derived ovarian cancer cells ex vivo. (A) Dual drug response assays for CLF + olaparib and CLF + rucaparib were performed in CLF-r OVAs (OVA-1, 2, 10, 11, 22, and 3) and in (C) CLF-nr OVAs (OVA-12, 15, 28, and 34). Each assay included three technical replicates, repeated independently three times (N=3), and was analyzed using the Loewe synergy and antagonism matrix model with Combenefit software. The larger numeral in each box of the synergy matrix represents the synergy score, with negative values indicating antagonism. Green boxes indicate non-significant synergy scores, while colored boxes correspond to statistically significant results based on the synergism/antagonism scale, determined using a one-sample t-test. (B and D) Combination index (CI) values across treatment panels were analyzed and presented in a table format. An average CI of 1 indicates an additive effect, CI < 1 represents synergy, and CI > 1 indicates antagonism.

### In CLF-Res cells, CLF does not downregulate CRM1 levels

Since CLF downregulated CRM1 levels in both CLF-r (Figure 1E and G) and-nr cells (Figure 1F and H), we surmised that CLF-resistant cells might have lost the ability to downregulate CRM1, resulting in a failure to retain CTSL in the nucleus. However, these cells might still traffic KPNB1-bound CTSL to the nucleus. Based on this premise, we predicted that these cells would respond to a combination therapy including a CRM1 inhibitor.

In our ongoing investigations, we discovered that certain cell lines, such as small cell lung carcinoma lines H1048 and H1882 and the endometrial cancer lines HEC1A, HEC1B, and RL95-2, were resistant to CLF-induced nCTSL, both alone and in combination with olaparib. Consistent with our hypothesis, we determined that CLF did not downregulate CRM1 in CLF-Res cells (Supplementary figure 9A Nuclear-Panel 2, Lane 2). However, treating RL95-2 cells with KPT8602 alone and in combination with either CLF and olaparib downregulated CRM1 expression (Supplementary figure 9A Nuclear Panel 2-Lanes 4, 6 and 7). Additionally, CLF (but not olaparib) retained the ability to import CTSL bound KPNB1 to the nucleus (Supplementary figure 9A Nuclear Panel 3, Lane 6). Consistent with this observation, CLF with KPT8602 and not KPT8602+olaparib combination resulted in the trafficking of nCTSL (Supplementary figure 9A Nuclear Panel 1, Lane 6). This finding suggests additional therapeutic options for sensitizing CLF-resistant cells using CRM1 inhibitors. Supporting this observation, clonogenic assays using RL-95-2 cells showed that KPT8602 synergized with CLF (Supplementary figure 9B-E).

Retesting the four CLF-Res samples, which previously showed antagonism with the CLF+olaparib combination, with the CRM1 inhibitor KPT8602 in combination with CLF or olaparib, revealed synergistic effects. Specifically, synergy was observed in OVA-9 and-16 with KPT8602+CLF combination and in OVA-29 and-30 with the KPT8602+olaparib combination, with CI values ranging from 0.4 to 0.7 (Supplementary figure 8C-F). These results provide further evidence that drug-induced nCTSL can sensitize cells to different combinations of the tested *drugs*.

### CLF synergizes with olaparib in FT33 cells transformed with TAg-Myc or shP53/CDK4^R24C^

To determine if the combination of CLF+olaparib is toxic to normal cells, we performed clonogenic assays in FT33 cells that are derived from non-diseased human fallopian tube secretory epithelial cells (FTSEC) by immortalization with human telomerase reverse transcriptase (hTERT) plus SV40 large T antigens (FT33TAg) in comparison to their counterparts that were further transformed with c-Myc or with sh-p53 and mutant CDK4 (CDK4^R24C^) [41]. Colony formation assays showed that FT33, FT33-shP53/CDK4^R24C^ and shPP2A-B56γ and FT33 c-Myc/TAg cells were resistant to CLF monotherapy up to 20 nM (Supplementary figure 10A). Clonogenic survival assays showed that FT33 cells treated with increasing concentration of olaparib + 5 nM CLF were resistant to this combination (Supplementary figure 10B). In contrast, under similar conditions, FT33-shP53-R24C that is transformed but not tumorigenic and FT33-TAg-Myc cells that are transformed and tumorigenic in nude mice were sensitized to olaparib by 5.0 nM CLF treatment (Supplementary figure 10C & D). Likewise, IF analysis showed that CLF and olaparib alone or in combination did not induce nuclear translocation of CTSL in FT33 cells, whereas CLF + olaparib promoted nCTSL in FT33TAg/Myc cells. Collectively, these data are consistent with the drug-induced nCTSL as both a driver and a mechanistically linked determinant of therapeutic response in transformed tumor cells and not in immortalized but nontransformed FTE cells.

### Nuclear translocation of CTSL during PDX treatment in vivo

The combination of CLF and olaparib or rucaparib significantly reduced tumor burden and ascites in mice bearing CLF-r PH610 (Figure 7A-C), while only the CLF+olaparib combination was effective in mice bearing CLF-nr PH747 (Figure 7F-H). Survival increased to 61 days in CLF-r and 56 days in CLF-nr mice with the CLF+olaparib treatment, compared to ∼40 days in the single-agent or CLF+rucaparib groups and controls (Figures 7D and I). Single-agent treatments in CLF-r mice also resulted in longer survival than untreated controls. Also, body conditioning score (BCS) study showed there was no significant body weight loss in CLF + Olaparib or CLF+ rucaparib combination groups compared to initial weight day 0 (Supplementary figure 11). Densitometric analysis of western blots from nuclear lysates of representative xenografts (n=3) showed a decrease in CRM1 with an increase in KPNB1 nuclear translocation accompanied by the presence of nCTSL, decreased 53BP1 and RAD51 expression, and an increase in γH2AX in the CLF+olaparib and CLF+rucaparib groups in CLF-r PH610 (Figure 7E). In CLF-nr PH747 xenografts, only the CLF+olaparib combination induced these changes (Figure 7J).

**Figure 7:**
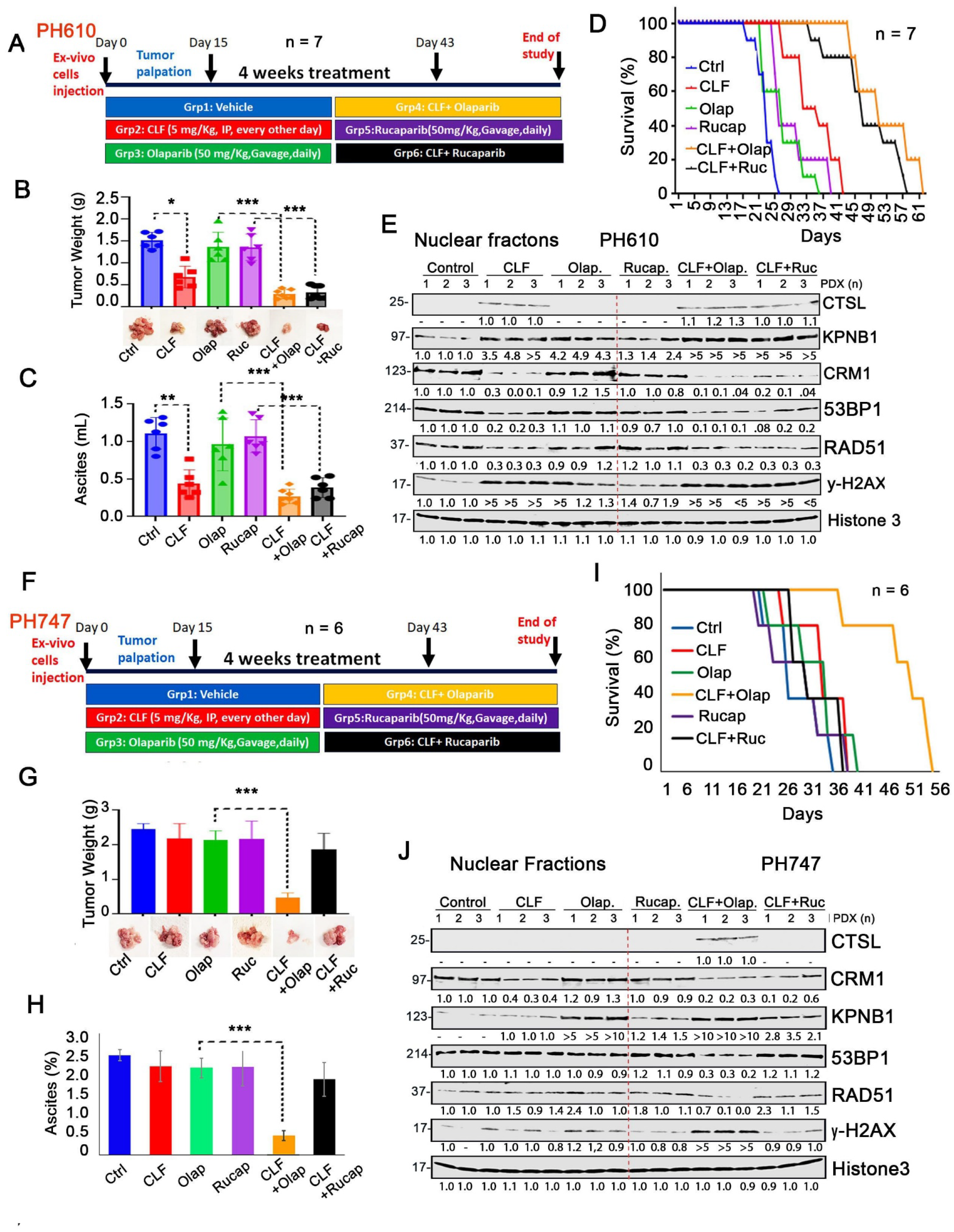
Evaluation of synergy upon treatment of CLF with the PARPis olaparib and rucaparib *in vivo* xenograft model of PH610 and PH747. (A, F) Schematic of the drug treatment regimen used in the in vivo xenograft models of PH610 (A) and PH747 (F). (B, G) Excised tumor weights comparing control and treated mouse cohorts in the PH610 (B) and PH747 (G) models. (C, H) Ascites volume collected from control and treated mice in the PH610 (C) and PH747 (H) xenograft models. (D, I) Overall survival analysis of control versus treated cohorts in the PH610 (D) and PH747 (I) models. Pair-wise log rank tests were performed and CLF + olaparib or rucaparib significantly improved survival vs control in PH610 (**p < 0.01: control vs combinations therapies, *p < 0.05: control vs CLF) and also only CLF + olaparib significantly enhanced survival vs control or monotherapies in PH747 model (**p < 0.01: Control vs CLF/Olaparib, **p < 0.01: CLF/rucaparib vs CLF/olaparib), whereas olaparib and rucaparib monotherapies were non-significant in either model. (E, J) Immunoblot analysis of nuclear-fractionated lysates showing levels of nCTSL, KPNB1, CRM1, and γH2AX in PH610 (E) and PH747 (J) xenografts, with Histone H3 serving as a nuclear loading control. Band intensities were quantified using ImageJ, normalized to the control, and fold changes are indicated below each panel.

### Efficacy of CLF and olaparib combination in KPCA syngeneic ovarian tumor model

The KPCA mouse model is a genetically defined, syngeneic ovarian cancer model derived from genetically engineered murine epithelial cells harboring oncogenic Kras and loss of the tumor suppressor Trp53, with overexpression of CCNE1 and Akt2 [33]. KPCA tumors are HR-proficient and resistant to PARP inhibition. We therefore evaluated the therapeutic efficacy of combined CLF and olaparib in this model. In KPCA-bearing mice (n = 10/group), the CLF/olaparib combination significantly reduced tumor growth by approximately two-fold compared with either monotherapy or vehicle control, identifying KPCA as a CLF-nr model (Supplementary figure. 12A-B). Consistent with the reduction in tumor growth, combination treatment significantly prolonged overall survival compared with CLF or olaparib monotherapy (Supplementary figure 12C), as demonstrated by pairwise log-rank analysis (p < 0.01). Only the CLF/olaparib combination, but neither monotherapy nor the control, induced nuclear CTSL, accompanied by decreased 53BP1 and RAD51 levels and increased γH2AX, as confirmed by western blot analysis of nuclear fractions isolated from KPCA tumors (Supplementary figure. 12D).

## Discussion

In our efforts to improve response to PARPis, we uncovered that nucleoside analog CLF, by promoting the translocation of cathepsin L to the nucleus, enhanced PARP inhibitor-induced cell death. Our data support the role of cathepsin L, a lysosomal cysteine protease that has functions beyond its conventional involvement in protein degradation in the cytoplasm. Cathepsin L has demonstrated contradictory roles in cancer by either worsening the transformed phenotype and as a tumor suppressor depending on the context and cellular environment. While the secreted CTSL isoform promotes migration and invasion [42, 43], regulated release of lysosomal CTSL contributes to crosstalk between lysosomal membrane permeabilization and mitochondrial membrane permeabilization leading to cell death [26, 44, 45]. Moreover, the nuclear isoform nCTSL has been implicated in Ras induced transformation [46], resistance to DNA damage inducing agents [47] and a bad prognosis [48]. In contrast to these latter reports, our data indicate that nCTSL promotes DNA damage, cell cycle arrest, cell death, and sensitivity to PARP inhibitors in cell line models *in vitro*, cultures of patient derived ovarian cancer *ex vivo*, and PDX models *in vivo*, providing support for a tumor suppressive role for drug-induced nCTSL (Figure 3-5 and 7). Consistent findings across cell lines, patient-derived ascites, and PDX models underscore the robustness and translational relevance of the mechanism.

Based on the report that CTSL, CUX1 and Snail all utilize importin β1 for nuclear import [35], we examined the role of KPNB1 in drug-induced nuclear CTSL import. Our data showed that one of the key molecular events for accumulation of CTSL in the nucleus involves the import of KPNB1-bound CTSL, which can be induced by CLF, in CLF-r cells and by olaparib in CLF-nr cells. This effect of olaparib appears to be independent of its ability to inhibit PARPs, as rucaparib lacks the ability to effect CTSL nuclear translocation in CLF-nr cells. Previous reports have separated the noncanonical functions of PARPis by showing that they bind different non-PARP proteins [49, 50]. An additional report indicates that the PARP inhibitor olaparib, but not talazoparib, acts as a mitochondrial Complex I inhibitor [51]. Also, studies by Kwarcinski et al., [52] showed how subtle changes in drug structure of dasatinib can influence binding selectivity and downstream effect. In support of KPNB1 bound cargo which can be induced by drugs, Xiao et.al, showed that following stimulation by TGF-β, the type I receptor phosphorylates the C-terminal region of Smad3, triggering a structural change that exposes its nuclear localization signal (NLS). This exposure allows importin-β to bind directly to Smad3, facilitating the nuclear import of the Smad3-Smad4 complex [53]. Moreover, our data indicate that CTSL preferentially utilizes its intrinsic nuclear localization signal (NLS) for nuclear trafficking, independent of the NLS present in Snail (supplementary figure 6)

Another critical alteration is CLF-induced downregulation of CRM1 expression, which aids in the nuclear retention of CTSL in both CLF-r and CLF-nr cells (Figure 1E-H). Moreover, our data also identified a subset of samples, which we termed CLF-Res cells, in which CLF fails to induce CRM1 downregulation and nuclear CTSL retention (supplementary figure 8). Interestingly, treating CLF-Res cells with the CRM1 inhibitor KPT8602 in combination with CLF or olaparib resulted in nuclear CTSL retention, induced DNA damage, and restored response to the CLF + PARPi combinations. Collectively, these findings suggest that CLF promotes the import of KPNB1-bound cargo into the nucleus in CLF-r cells but requires interruption of CRM1 function, e.g., through olaparib-induced downregulation of CRM1 in CLF-nr cells or treatment with KPT8602 CLF-Res cells, for nuclear retention (Supplementary figure 9).

It is important to note that CLF has dual function in CLF-r cells: 1) facilitating import of CTSL-bound KPNB1 into the nucleus and 2) attenuating CRM1 levels to retain CTSL in the nucleus. This dual function allows CLF to synergize with both olaparib and rucaparib (potentially with other PARP inhibitors) in CLF-r cells. In contrast, in CLF-nr cells, while CLF downregulates CRM1 expression, olaparib—but not rucaparib—facilitates the import of CTSL-bound KPNB1 into the nucleus (Figure 1C-H, Figure 2A-B). We confirmed the differential binding of CLF and olaparib to their target KPNB1 using the CETSA assay in CLF-r versus CLF-nr cells respectively, providing evidence for the differential drug response in the import of KPNB1-bound CTSL to the nucleus (Figure 2C-I). Taken together, our data identify KPNB1-dependent import and CRM1-dependent retention as key regulatory nodes, establishing nuclear trafficking as a mechanistic layer controlling DNA repair capacity. Consequently, these results present a promising opportunity for using a CRM1 inhibitor with CLF or olaparib as a novel combination therapy for these highly resistant cells.

At the same time, the mechanism of CLF-induced CRM1 downregulation is not currently understood. It is also unknown why CLF has lost the ability to downregulate CRM1 in CLF-Res cells. We observed in RNA seq (data not shown), no changes in XPO1 transcript abundance following treatment, indicating that CRM1 downregulation is unlikely to occur through transcriptional regulation and is instead consistent with a post-transcriptional mechanism. Additionally, based on available TCGA data, CRM1 (XPO1) is not recurrently mutated in ovarian cancer. Based on results showing that CLF can act as a demethylating agent in breast cancer [8, 10], it is possible that CLF-induced demethylation of the *CRM1* promoter may lead to upregulated levels CRM1 in CLF-Res cells described in the present study. Consistent with this possibility, studies by Yamanashi et al. suggest that histone H3 and H4 acetylation are significantly decreased in CLF-Res cells, and a melatonin induced increase in acetylation may overcome CLF resistance in leukemia cell lines [54].

Our further studies showed that the CLF+olaparib combination induced nCTSL in both CLF-r and CLF-nr cells as well as synergistic antiproliferative effects in over 70% of clinical ovarian cancer specimens. Similarly, in CLF-Res samples, KPT8602 in combination with CLF or olaparib promoted nCTSL and synergy in 33% of the samples (Figure 6). This unique observation that all 14 OVA samples, drug resistant cell lines (HeyA8MDR, PEO1/OLTRC4, C13, OVCAR 8), and *ex vivo* cultures of PARPi resistant PDXs (PH610, PH747) show synergy when nCTSL was promoted indicate a tight association between drug-induced nCTSL and response, raising the possibility that therapy-induced nCTSL could potentially be evaluated in cultures of patient-derived tumor samples *ex vivo* to identify effective therapies for individual ovarian cancers. Further studies are required to evaluate this possibility.

We find that CLF selectively cooperates with olaparib—but not rucaparib, niraparib, or veliparib—to drive robust nuclear CTSL (nCTSL) translocation, functional heterogeneity among PARP inhibitors and that PARP inhibitors are not functionally equivalent in engaging CTSL-dependent DNA repair responses. While our current data support enhanced olaparib–KPNB1 interactions, recent evidence that rucaparib and niraparib undergo lysosomal sequestration suggests an alternative mechanism [55]. Lysosomal trapping may restrict nuclear drug availability and attenuate signaling required for CTSL trafficking. In contrast, olaparib sustains nuclear PARP inhibition, promoting accumulation of nCTSL and activation of downstream DNA damage and repair signaling.

The role of nCTSL defined in ovarian cancer in the present study is somewhat different from the previously reported roles of nCTSL in breast cancer. Initially, Gortsky et al. [17] reported that growth arrested cells transcriptionally downregulated BRCA1 (not BRCA2) and activated nCTSL-mediated degradation of 53BP1, facilitating homologous recombination repair (HRR) [47] and enabling triple-negative breast cancer cells to survive in the absence of BRCA1 [56]. In a follow-up study, Croke et al. [57] demonstrated that nCTSL-mediated 53BP1 degradation is a natural regulatory mechanism that balances non-homologous end joining (NHEJ) versus HRR repair in non-cycling cells: When cell reenter the cell cycle, BRCA1 expression is upregulated and CTSL is downregulated, restoring 53BP1 and leading to increased genomic instability, particularly in response to radiation and PARP inhibitors. Saurez et al., [47] reported that A-type lamin loss favors CTSL-mediated degradation of 53BP1and treatment of WT and Lmna−/− MEFs with proteosome inhibitor MG-132 or the CTSL inhibitor Z-FY-CHO, stabilized 53BP1 implicating both pathways in it’s degradation. Our data using rCTSL also showed that CTSL mediated degradation of 53BP1 and RAD51 were rescued in the presence of CTSL specific inhibitor (Figure 3G). In contrast to these previous studies, our data show that drug-induced nCTSL also downregulates RAD51, thus inhibiting HRR. Additionally, the effects of drug-induced nCTSL reported in this study are independent of BRCA1 status. Thus, although prior studies have described nuclear CTSL in limited contexts, our work identifies a drug-inducible, trafficking-dependent trafficking axis that governs its nuclear localization. Importantly, we demonstrate that nCTSL simultaneously targets both 53BP1 and RAD51, establishing a previously unrecognized mechanism of homologous recombination collapse. Consistent with these findings, CTSL depletion, pharmacologic inhibition, and recombinant protein assays collectively establish a causal role for CTSL in DNA repair disruption and therapeutic response.

Previous work by Tholen et al., [58] has argued that downstream AUG usage in murine CTSL is not physiologically relevant for CTSL, proposing that out-of-frame AUGs prevent efficient translation of truncated isoforms and that nuclear CTSL may therefore be artifactual. While murine CTSL does contain in-frame downstream AUGs, these studies examined constitutive translation in the absence of stress and did not account for transcript-specific or context-dependent initiation. In human cells, we show that CTSL translation is dynamically regulated, with stress conditions favoring production of an M56-initiated isoform that undergoes robust nuclear translocation in vitro and in primary tumors. These findings indicate that nuclear CTSL reflects regulated, stress-dependent translation rather than nonspecific downstream initiation.

In summary, while previous data suggest that effects of nCTSL might be highly context-dependent, our results suggest that the drug-induced induction of nCTSL is tightly linked to downregulation of key DNA damage response protein in multiple pathways followed by induction of cell death (Figure 8), raising the possibility that nCTSL might be further used to identify effective treatment options. Moreover, our results identify CLF as a nucleoside analog that enhances olaparib cytotoxicity in CLF-r and CLF-nr cells and the CRM1 inhibitor KPT8602 as an agent that overcomes CLF resistance to facilitate killing by olaparib or CLF, identifying two new combinations that warrant further investigation in the effort to overcome PARPi resistance. nCTSL is induced under defined stress conditions and observed across primary samples and in vivo models, supporting its regulated and physiologically relevant production. While our data establishes a strong mechanistic and functional association between nCTSL and therapeutic response across multiple model systems, prospective clinical validation will be required to determine its utility as a predictive functional biomarker in patients.

**Figure 8.**
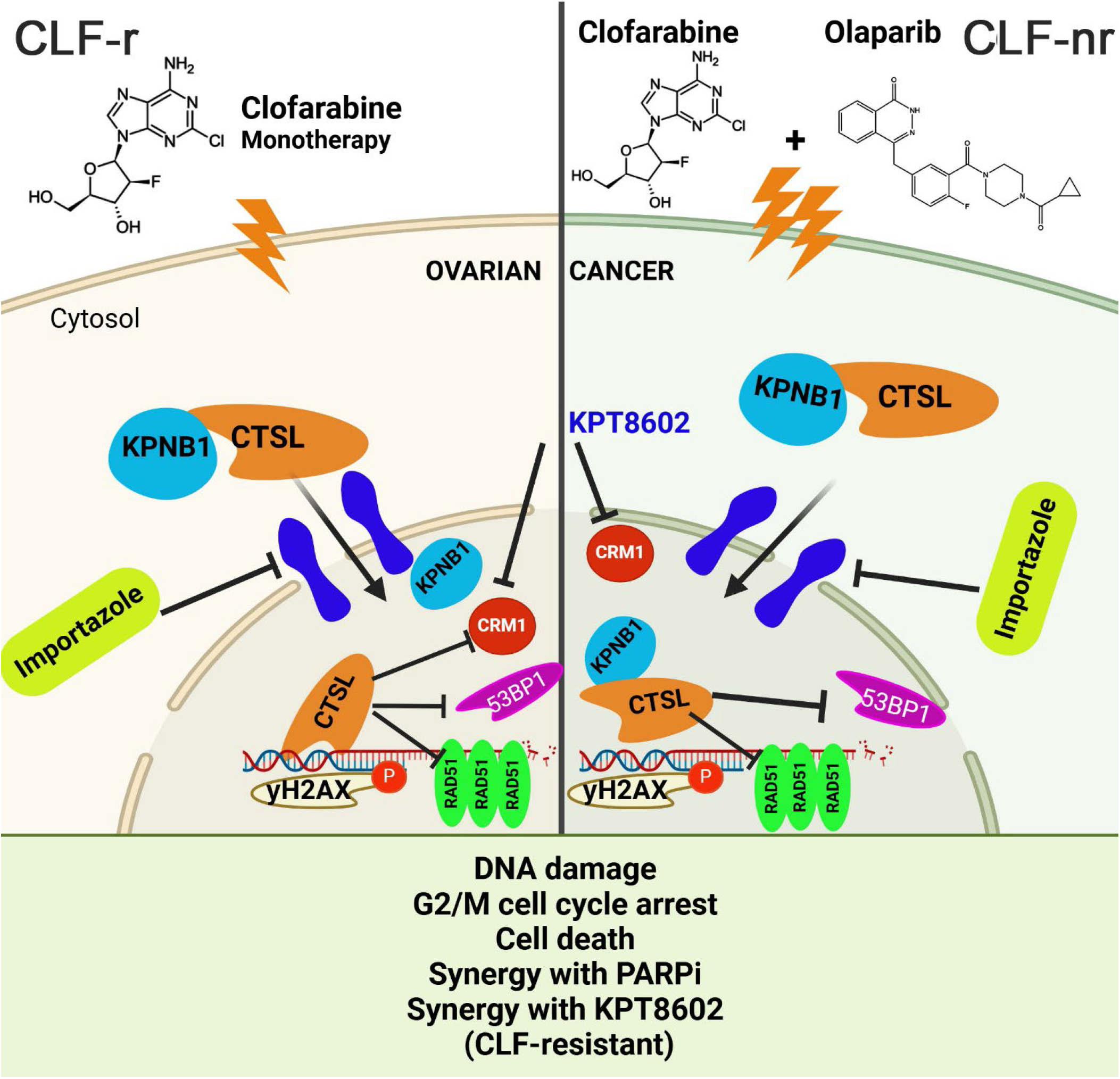
A drug-inducible KPNB1–CRM1–CTSL trafficking axis enforces DNA repair collapse and PARP inhibitor sensitivity. We propose that therapeutic stress activates a KPNB1–CRM1–CTSL nuclear trafficking axis that reprograms the DNA damage response. Clofarabine (CLF) promotes KPNB1-dependent nuclear import of CTSL, while CRM1 downregulation enables its nuclear retention, resulting in accumulation of nuclear CTSL (nCTSL). Nuclear CTSL degrades 53BP1 and RAD51, driving DNA repair collapse, DNA damage accumulation and caspase independent cell death (BioRender, https://biorender.com). Abbreviations; CLF-- clofarabine, R-responsive, NR-Nonresponsive, CTSL-cathepsin L, CRM1-chromosomal region maintenance 1, KPNB1-karyopherin β1, 53BP1-p53-binding protein 1, RAD51-RAD51 recombinase, ϒH2AX-gamma H2A histone family member X, Olaparib-PARP inhibitor, Importazole-KPNB1 inhibitor, KPT8602-CRM1 inhibitor.

## Acknowledgements

We would also like to acknowledge the Mayo Ovarian SPORE for collection of OC ascites samples from patients. In addition, we thank Dr. Larry Karnitz (Mayo Clinic Rochester, MN) for OVCAR8 DR-GFP cells, Drs. Amy Skubitz and Kristin Boylan for cryopreserved patient-derived ascites samples from the University of Minnesota under an IRB-approved protocol, and Dr. Ronny Drapkin (University of Pennsylvania Perelman School of Medicine) for the FT33(hTERT/TAg), FT33-shP53/CDK4^R24C^ and shPP2A-B56γ and FT33 c-Myc/TAg cells. We gratefully acknowledge contributions of the Mayo Clinic Cancer Center (P50 CA015803)-supported Pathology Research and Microscopy & Cell Analysis Cores.

## Contributions

Conceptualization: VS and PT. Methodology & Investigation: PT, LJ, AZ, SR., XW, XH and JLV. Data assessment and verification: LJ, PT, UR, JS, SMMA, AO, NK, JW, JC, SHK, JNB and VS. Supervision: VS. Writing—original draft: VS and PT. Writing—review & editing: JC, UR, SHK and VS. Correspondence and requests for materials should be addressed to VS.

## Clinical trial number

not applicable

## Data availability

Data is available within the Article or Supplementary Information.

## Ethics approval and consent to participate

All procedures involving animal were in accordance with the ethical standards of the ethics committee of Mayo Clinic Animal Care and Use Committee (IACUC number#A00006931, A00007937)). Clinical samples from patients were obtained at Mayo Clinic (IRB-approved protocol; 1288-03) and in association with the University of Minnesota Cancer Center Tissue Procurement Facility (IRB approval; 0702E01841).

