## Supplemental files for "Drug-induced nuclear cathepsin L (nCTSL) defines a targetable DNA damage response axis and PARP inhibitor sensitivity in ovarian cancer"

**Supplemental information**

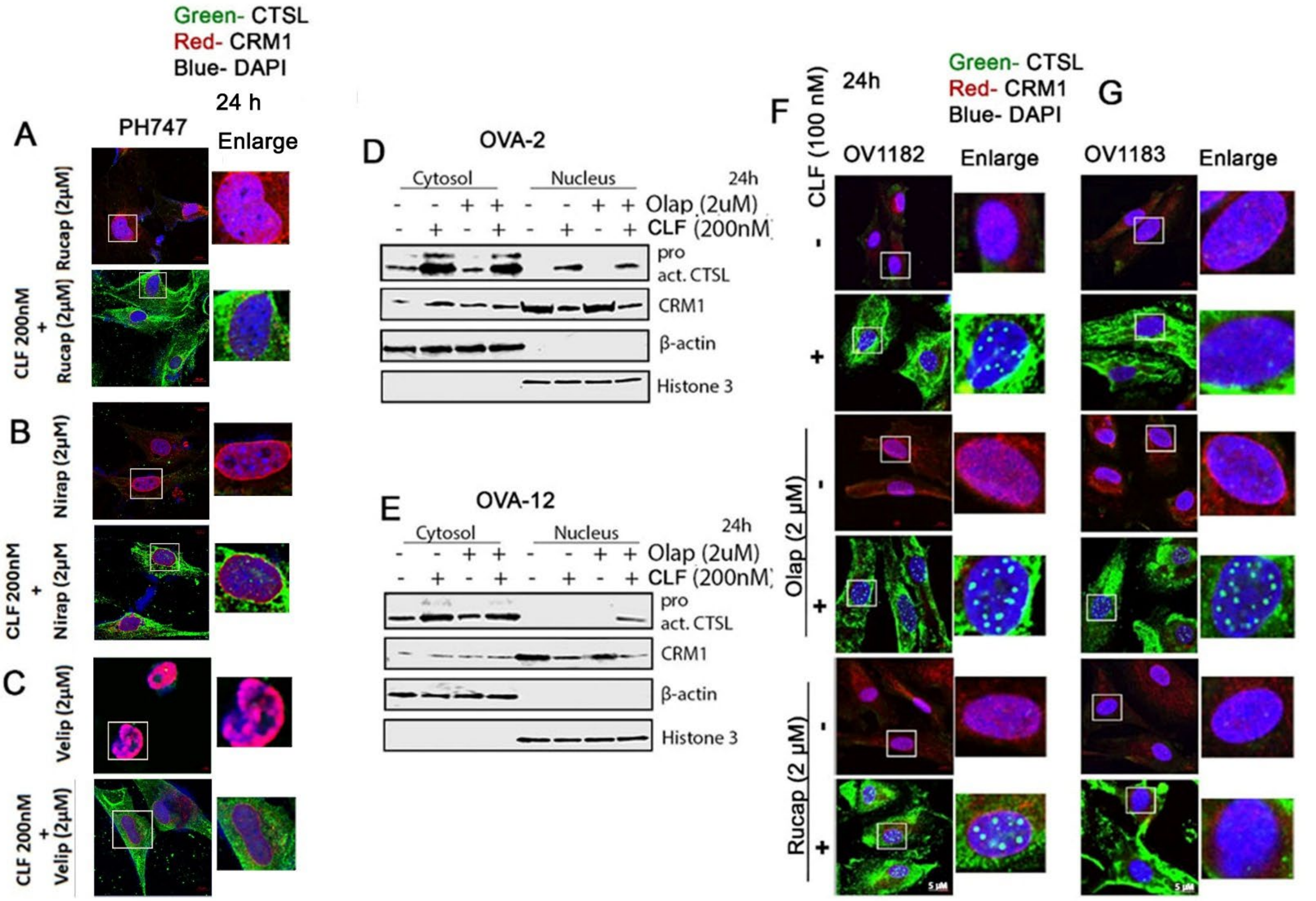


**Supplementary Figure 1: Nuclear translocation of CTSL after CLF or CLF + olaparib treatment.** Representative images of the immunofluorescence staining of CTSL (green) and CRM1 (red) are shown in PH747 cells upon treatment with CLF monotherapy or in combination with (A) rucaparib, (B) niraparib and (C) veliparib for 24 hours. (40X magnification, scale bar (5µm). DAPI was used to stain the nucleus. Immunoblot analysis of cell fractionated lysates in (D) OVA-2 and (E) OVA-12 for CTSL and CRM1. β-actin and Histone H3 were used as endogenous controls for the cytoplasmic and nuclear fractions, respectively. (F and G) Ex vivo cultures of OV1182 and OV1183 after treatment with CLF, either as a single agent or in combination with the PARP1 inhibitors olaparib and rucaparib, at the indicated concentrations for 24 hours. Representative images show DAPI staining the nucleus (40X magnification, scale bar: 10 μm).

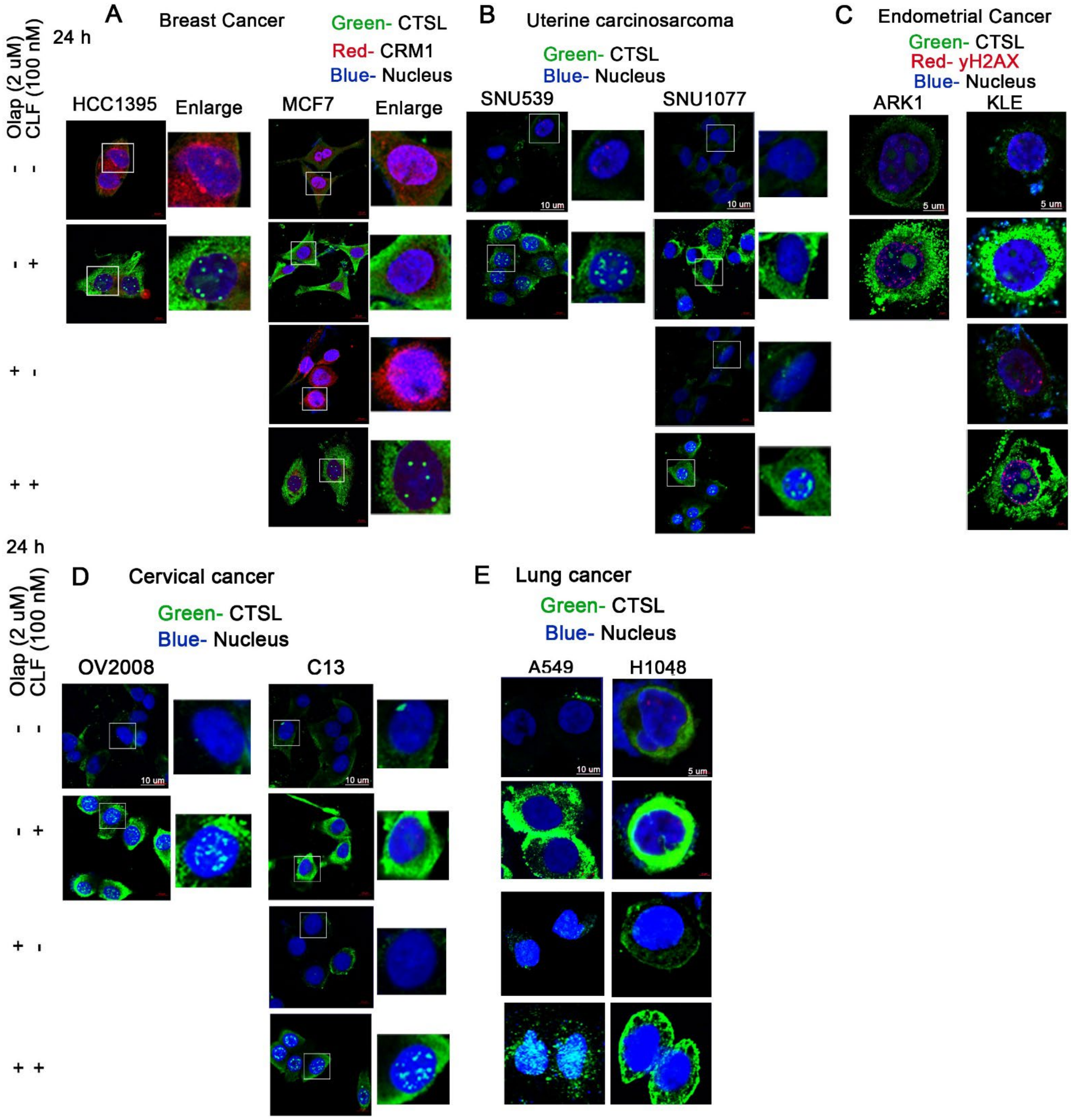


**Supplementary Figure 2: Nuclear presence of CTSL after CLF or CLF + olaparib treatment on various tumor types.** (A) Breast cancer (B) uterine carcinosarcoma (C) endometrial cancer (D) cervical cancer and (E) lung cancer cells were treated with CLF and olaparib at indicated concentrations for 24 hours and immunofluorescence (IF) analysis was performed. Representative images of the IF staining of CTSL, CRM1 or yH2AX and DAPI are shown (40X magnification, DAPI was used to stain the nucleus.

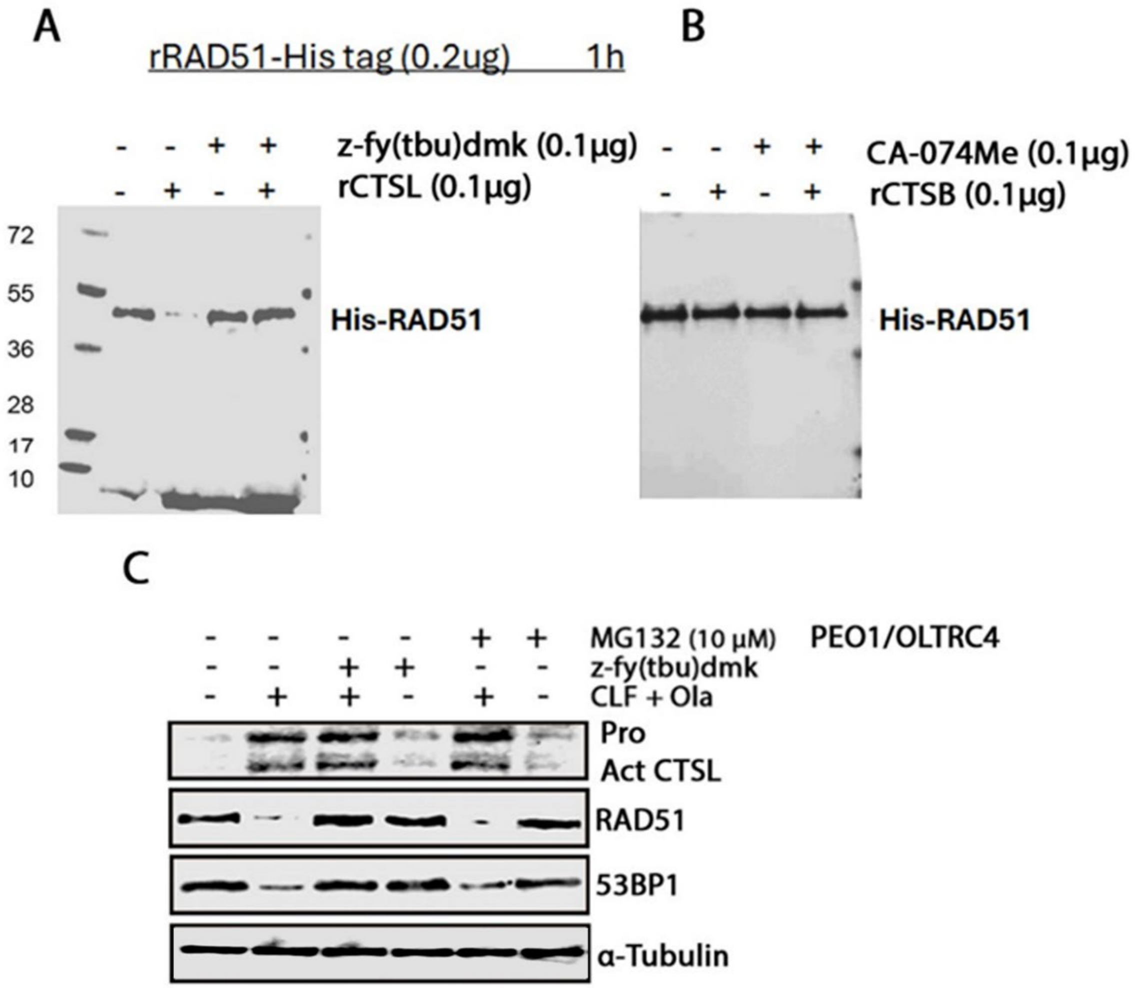


**Supplementary figure 3: nCTSL degrades both RAD51 and 53BP1 independent of proteasome.** (A and B)His-tagged recombinant RAD51 (0.2 μg) incubated with recombinant CTSL or CTSB at indicated concentration in assay buffer for 1 h and then, western blots were performed using anti-His antibody (C) Western blot analysis was performed using lysates of PEO1/OLTRC4 cells treated with CLF (100 nM) and olaparib (2 µM) in presence of CTSL inhibitor z-fy(tbu)dmk (5 µM) or proteasome inhibitor MG132 (10µM) for 24h. Blots were probed with the CTSL, RAD51 and 53BP1 antibodies, with α-tubulin used as endogenous control.

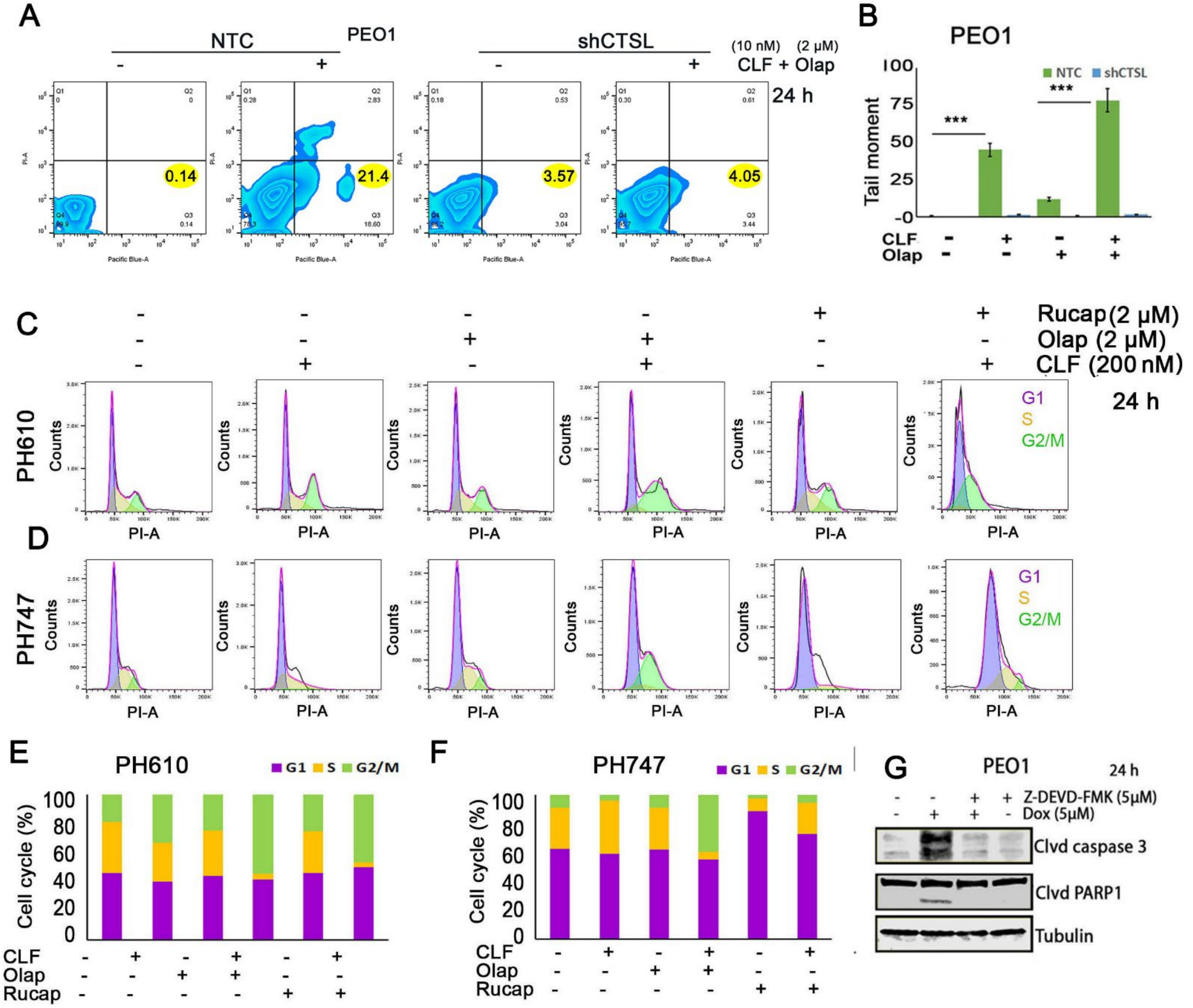

**Supplementary figure 4: CTSL knockdown restores CLF induced DNA damage in OC cells and CLF in combination with olaparib arrests the cell cycle at G2M in PDX cells**

(A) Annexin V/PI staining of PEO1-NTC and sh2-CTSL cells treated with CLF (10 nM) and olaparib(2 μM) for 24 h. (B) The alkaline COMET assay was also conducted in PEO1-NTC and CTSL knockdown (sh2-CTSL) cells following treatment with the indicated concentrations of CLF and olaparib, either alone or in combination, for 24 hours. For all cases, quantification of the tail moment is shown (***$p < 0.001$ vs. control). (C-D) Cell cycle analysis was conducted in PH610 and PH747 cells using PI staining and flow cytometry after treatment with the indicated concentrations of CLF, olaparib, and rucaparib, either alone or in combination, *ex vivo* for 24 hours. (E-F) The percentage of cells in each cell cycle phase was quantified and plotted for PH610 and PH747 cells, respectively. (G) Western blot analysis of whole-cell lysates from PEO1 cells treated with doxorubicin (5 μM) with or without Z-DEVD-FMK (5 μM) for 24 hours. Blots were probed for cleaved caspase-3 and PARP1, with tubulin as a loading control.

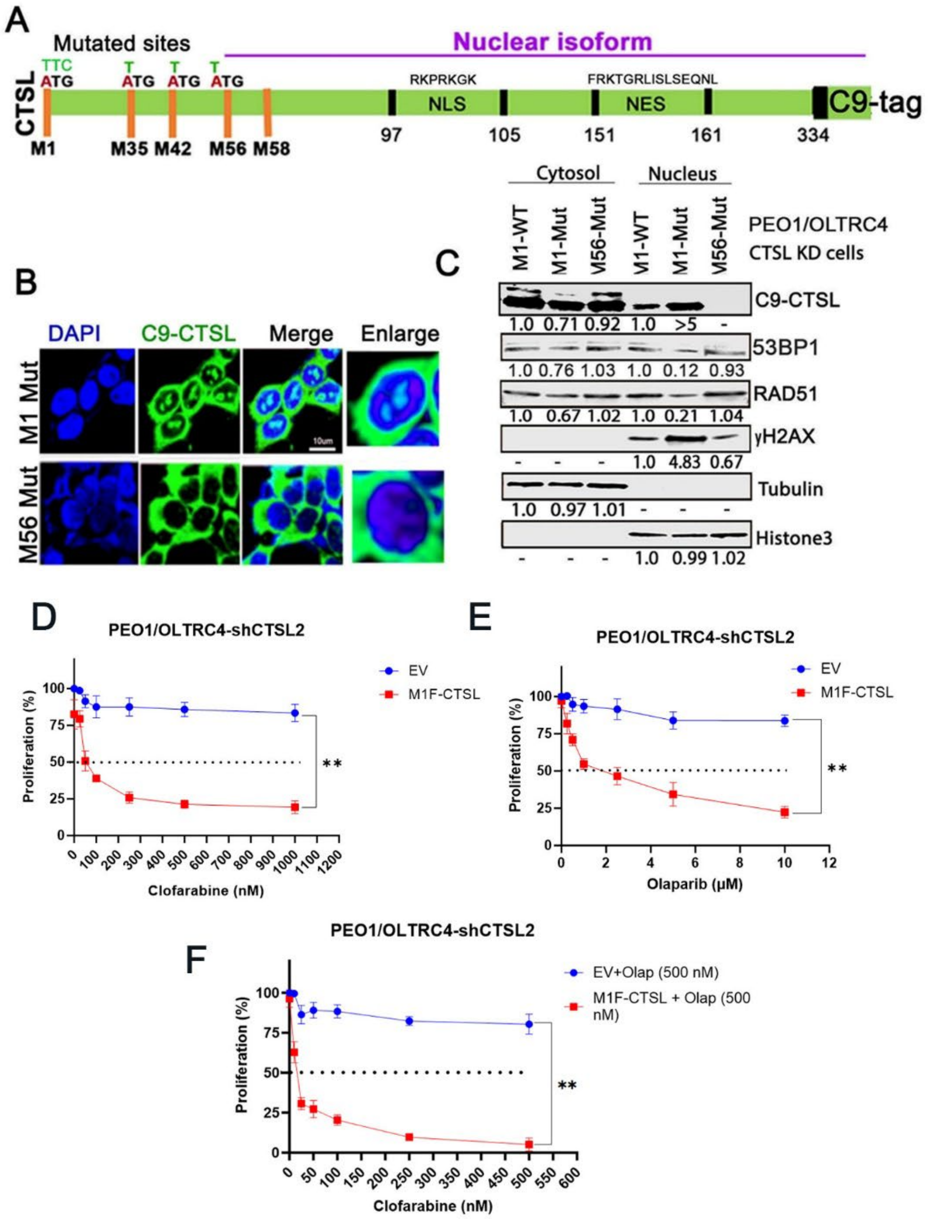


**Supplementary figure 5: nCTSL is specifically involved in the DNA damage response.** (A) Schematic representation of the C9-tagged CTSL construct, highlighting the alternative start codons, nuclear localization signal (NLS), and nuclear export sequence (NES) motifs. (B) Immunofluorescence analysis of C9-tagged M1 and M56 mutant constructs in PEO1/OLTRC4sh2 CTSL cells, detected with an anti-C9 antibody. (C) Western blot analysis of cytoplasmic and nuclear fractions was performed in PEO1/OLTRC4sh2 CTSL cells transfected with the M1 WT, M1-Mut, and M56-Mut nCTSL isoform constructs. Blots were probed with the indicated antibodies, with tubulin and Histone H3 used as endogenous controls for the cytoplasmic and nuclear fractions, respectively. PEO1/OLTRC4-shCTSLL-2 cells transfected with vector and M1F-CTSL and treated with CLF and olaparib at indicated concentrations for 24 hours. CCK8 proliferation assay was performed and proliferation rate was plotted for (D) CLF alone, (E) olaparib alone and (F) CLF and olaparib combinations. Data represent mean ± SD; $^{**}p < 0.001$ vs empty vector (EV).

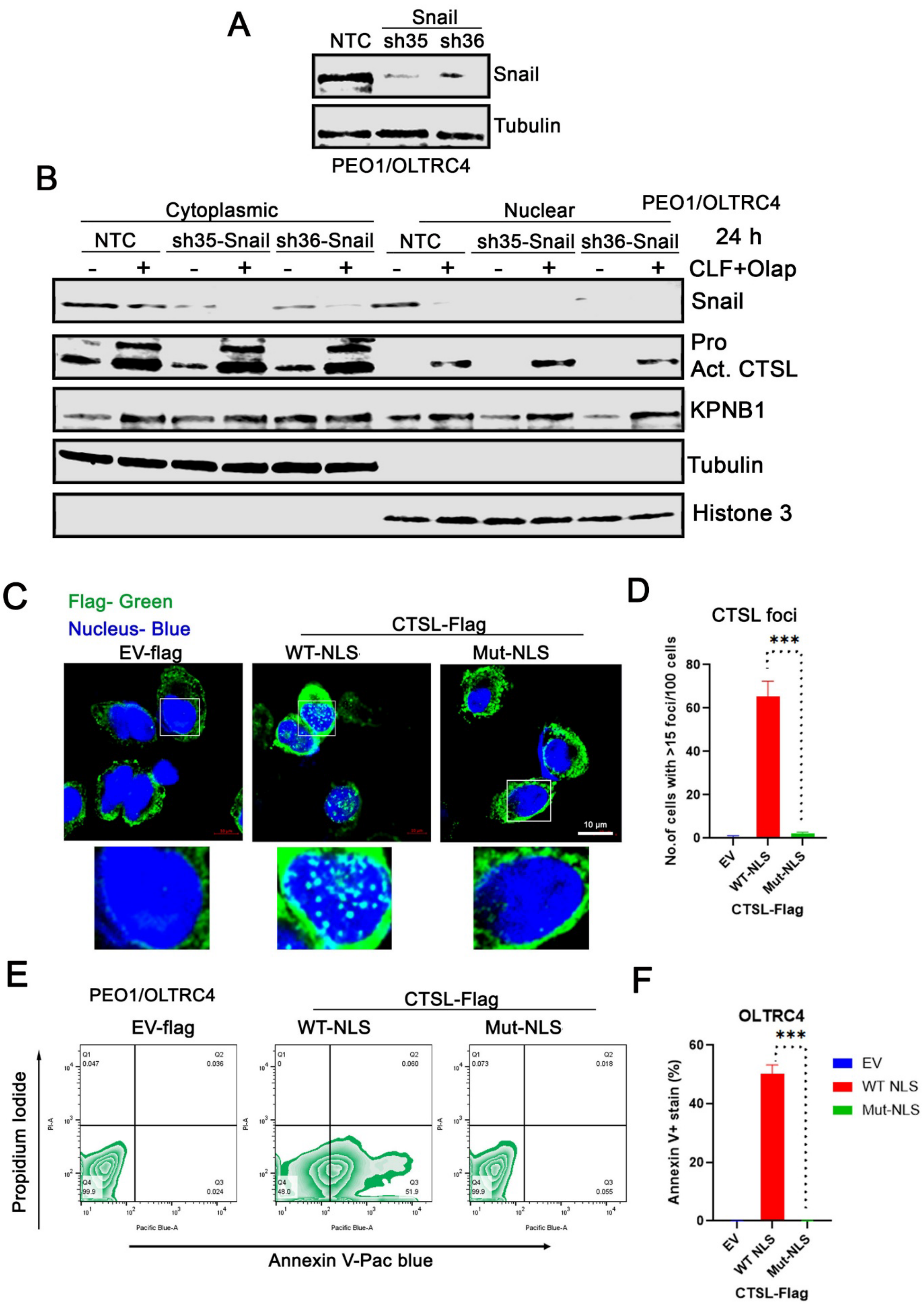
A
Snail
NTC
sh35
sh36
Snail
Tubulin
PEO1/OLTRC4
B
Cytoplasmic
Nuclear
PEO1/OLTRC4
NTC
sh35-Snail
sh36-Snail
NTC
sh35-Snail
sh36-Snail
24 h
CLF+Olap
Snail
Pro
Act. CTSL
KPNB1
Tubulin
Histone 3
C
Flag- Green
Nucleus- Blue
CTSL-Flag
EV-flag
WT-NLS
Mut-NLS
10 µm
D
CTSL foci
***
No.of cells with >15 foci/100 cells
EV
WT-NLS
Mut-NLS
CTSL-Flag
E
PEO1/OLTRC4
CTSL-Flag
EV-flag
WT-NLS
Mut-NLS
Propidium Iodide
Annexin V-Pac blue
Pacific Blue-A
PI-A
Q1 0.047
Q2 0.036
Q4 99.9
Q3 0.024
Q1 0
Q2 0.060
Q4 48.0
Q3 51.9
Q1 0.073
Q2 0.018
Q4 99.9
Q3 0.055
F
OLTRC4
***
Annexin V+ stain (%)
EV
WT NLS
Mut-NLS
CTSL-Flag

**Supplementary figure 6: A**) Western blot confirming Snai1 (Snail) knockdown in PEO1 and OLTRC4 cells using shSnail compared with non-targeting control (NTC). (B) Immunoblot analysis of fractionated lysates from PEO1/OLTRC4 NTC and Snail-knockdown cells treated with CLF (100 nM) and olaparib (2 μM) for 24 h. Blots were probed with antibodies against Snail, CTSL, and KPNB1, with α-tubulin and histone H3 serving as cytosolic and nuclear loading controls, respectively. (C) Representative immunofluorescence images of CTSL-Flag (green) and nuclei (DAPI, blue) in PEO1 and OLTRC4 cells transfected with empty vector, NLS-wild-type CTSL-Flag, or NLS-mutant CTSL-Flag constructs (40× magnification; scale bar, 5 μm). (D) Quantification of cells containing >15 CTSL nuclear foci per 100 cells. (E–F) Annexin V/PI apoptosis assays in PEO1 and OLTRC4 cells expressing empty vector, NLS-wild-type CTSL, or NLS-mutant CTSL constructs. Data represent mean ± SEM; ***p < 0.001 vs. control or empty vector (EV).

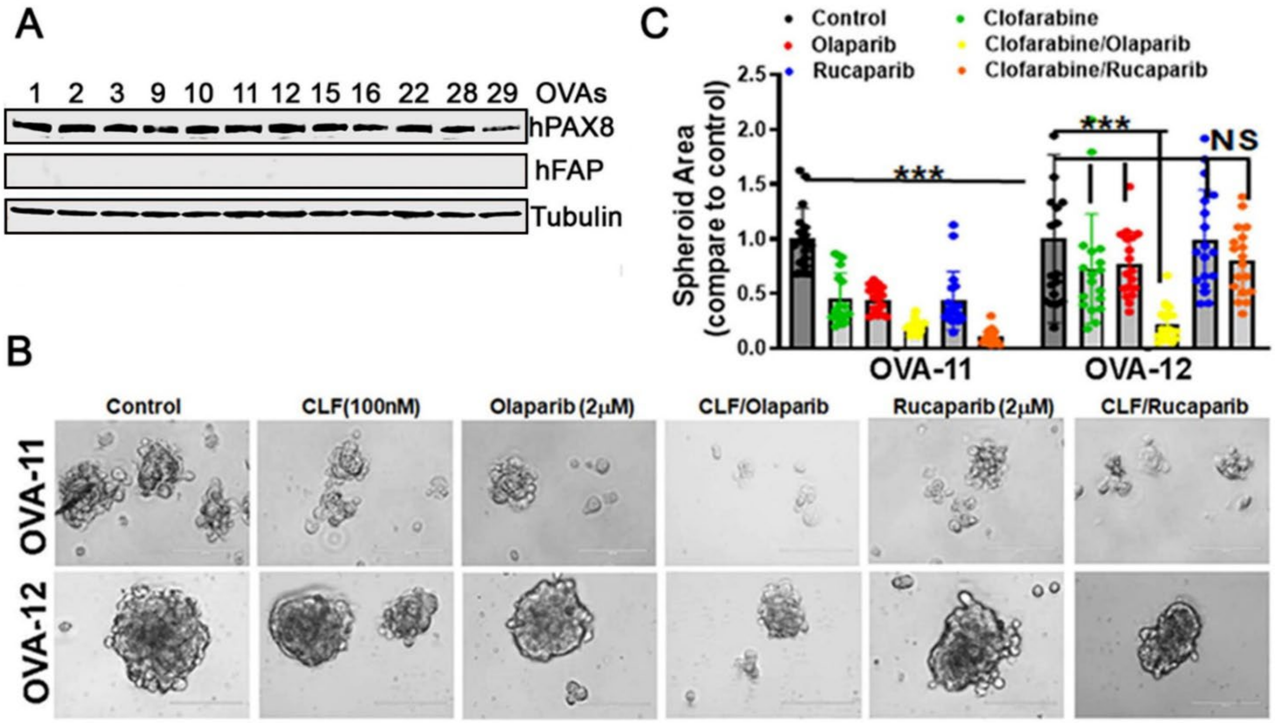

**Supplementary figure 7:** (A) Immunoblot analysis of epithelial cell marker PAX8 and fibroblast specific marker hFAP in OVAs cells. Tubulin is used as a loading control. (B) Representative images of spheroid forming ability of patients derived ovarian cancer ascitic cells OVA-11 and OVA-12 and their response to CLF and mentioned PARP inhibitors. (C) Bar graphs show the area of spheroids in untreated and treatment groups.

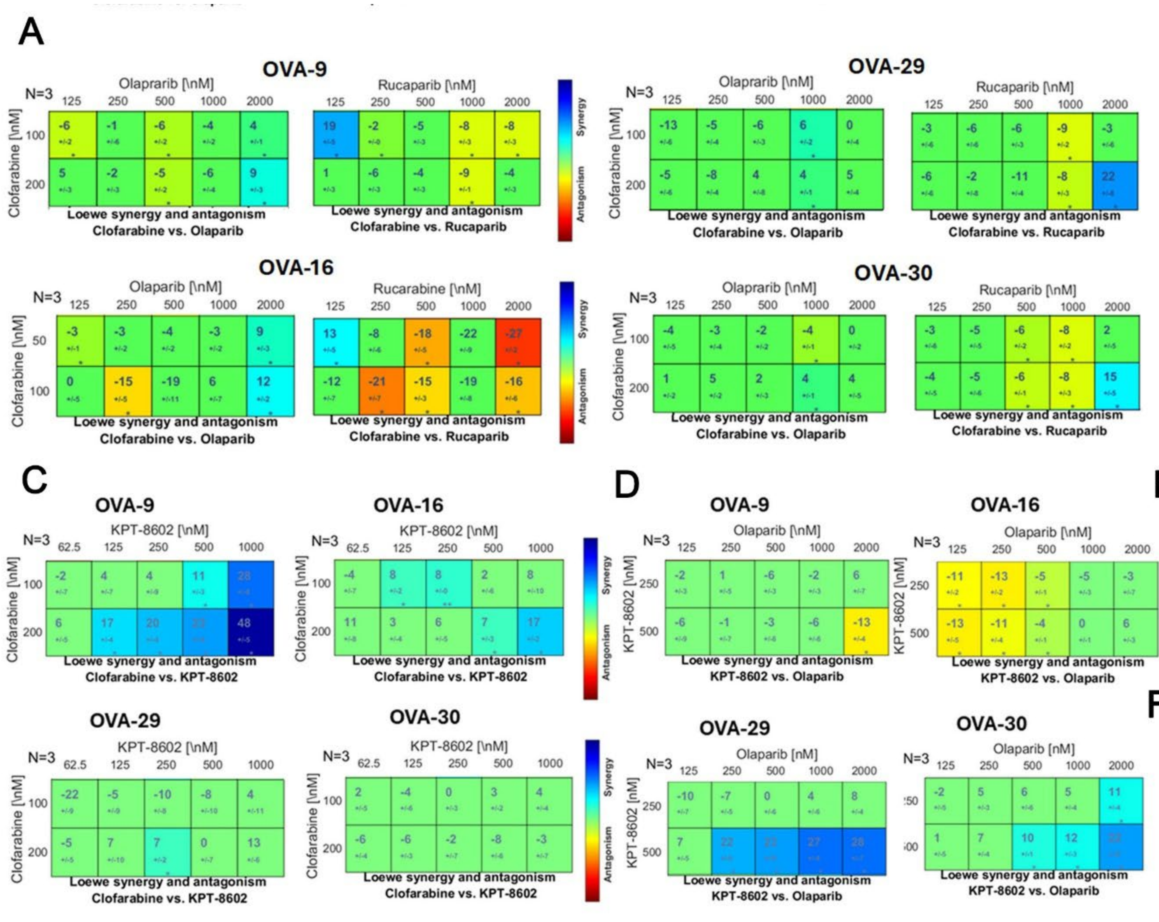


B

| | CLF (nM) | CI Value | |
|---|---|---|---|
| | | Olap | Rucap |
| OVA-9 | 100 | 2.017 | >10 |
| | 200 | >10 | >10 |
| OVA-16 | 100 | 4.912 | NaN |
| | 200 | >10 | NaA |
| OVA-29 | 100 | NaN | NaN |
| | 200 | NaN | NaN |
| OVA-30 | 100 | 1.613 | 2.564 |
| | 200 | 1.062 | 5.084 |

E

| | CLF (nM) | CI Value |
|---|---|---|
| | | KPT-8602 |
| OVA-9 | 100 | 0.156 |
| | 200 | 0.123 |
| OVA-16 | 100 | 0.728 |
| | 200 | 0.647 |
| OVA-29 | 100 | NaN |
| | 200 | NaN |
| OVA-30 | 100 | 2.94 |
| | 200 | 1.196 |

F

| | KPT-8602 (nM) | CI Value |
|---|---|---|
| | | Olap |
| OVA-9 | 250 | 1.048 |
| | 500 | 2.1 |
| OVA-16 | 250 | >10 |
| | 250 | >10 |
| OVA-29 | 500 | 1.416 |
| | 500 | 0.026 |
| OVA-30 | 250 | 0.597 |
| | 500 | 0.399 |

**Supplementary figure 8: Evaluation of synergy upon treatment of CLF with olaparib and rucaparib respectively in the ascites-derived ovarian cancer cells ex vivo.** (A) Dual drug response assays for CLF + olaparib and CLF + rucaparib were performed in CLF-Res resistant OVAs (OVA-9, 16, 29, and 30). Each assay included three technical replicates, repeated independently three times (N=3), and was analyzed using the Loewe synergy and antagonism matrix model with Combenefit software. (C and D) Additional drug response assays for CLF + KPT8602 and olaparib + KPT8602, respectively were conducted in CLF-Res-resistant OVAs

(OVA-9, 16, 29, and 30), analyzed, and represented similarly. The larger numeral in each box of the synergy matrix represents the synergy score, with negative values indicating antagonism. (B, E, F) Combination index (CI) values across treatment panels were analyzed and presented. An average CI of 1 indicates an additive effect, CI < 1 represents synergy, and CI > 1 indicates antagonism.

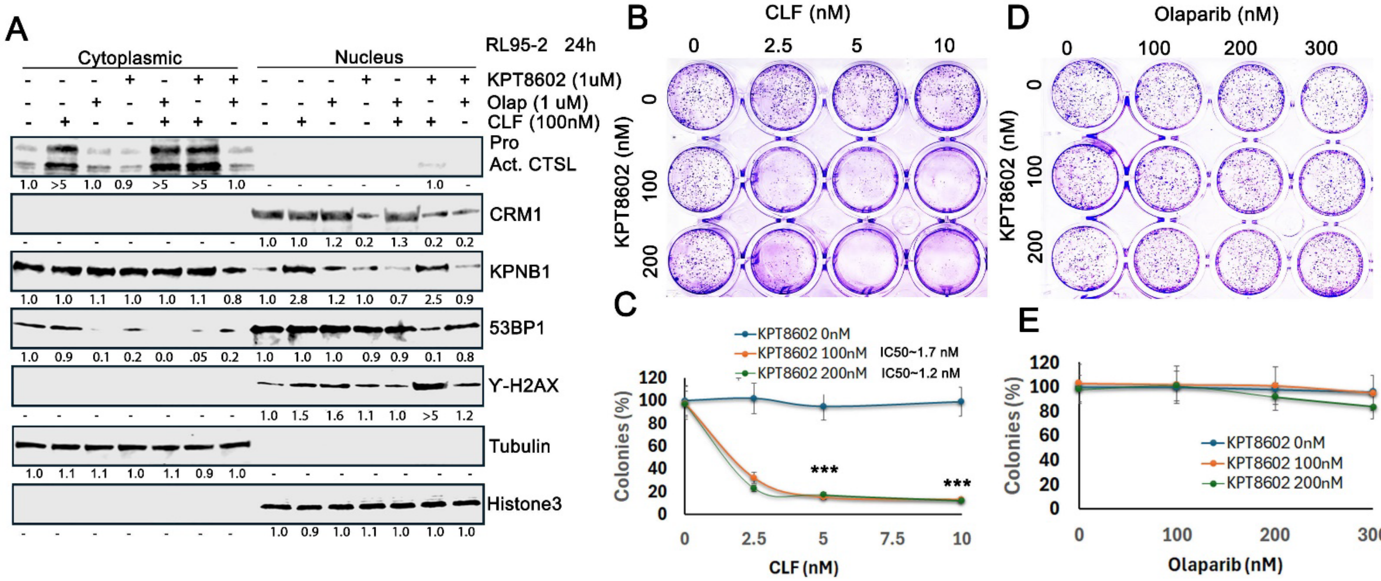


**Supplementary figure 9: KPT8602 sensitizes CLF resistant RL-95-2 to CLF treatment by downregulating CRM1.** (A) Western blot analysis of cytoplasmic and nuclear fractions from CLF-resistant RL-95-2 cells was performed to assess CTSL, CRM1, KPNB1, and the nuclear control Histone H3 levels following treatment with the indicated drugs for 24 hours. Fold changes were calculated using ImageJ software, normalized, and displayed below each panel. (B and D) A clonogenic assay was conducted in RL-95-2 cells treated with the indicated concentrations of CLF alone or in combination with olaparib and KPT8602. (C and E) Quantification of the clonogenic assay was represented as colonies (% of the control) in the RL95-2 cells. Error bars represent ± SEM from triplicates of a single assay (n=3).

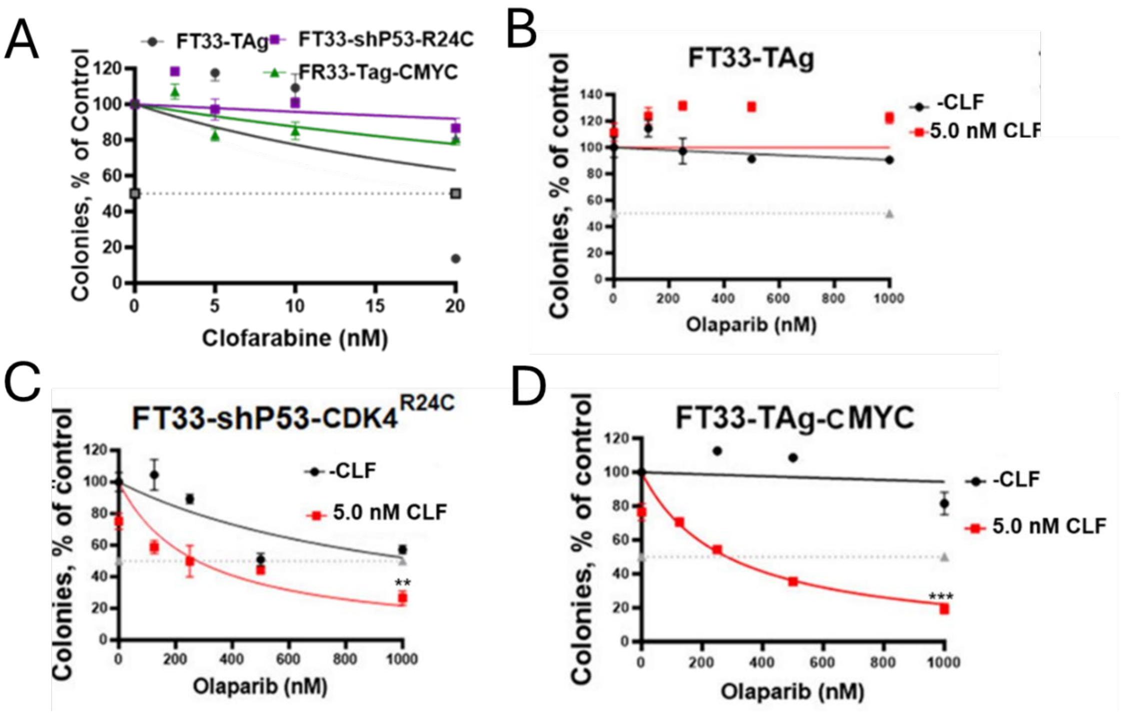


**Supplementary figure 10:** Cytotoxicity assessment by colony forming assay upon treatment with clofarabine alone in FT33-TAg, FT33-shP53-CDK4(R24C) and FT33-TAg-cMYC in (A) and with – and + 5 nM CLF in combination with olaparib at indicated concentration in (B) FT33-TAg, (C) FT33-shP53-CDK4(R24C) and (D) FT33-TAg-cMYC. Error bars represent ± SEM from triplicates of a single assay (n=3).

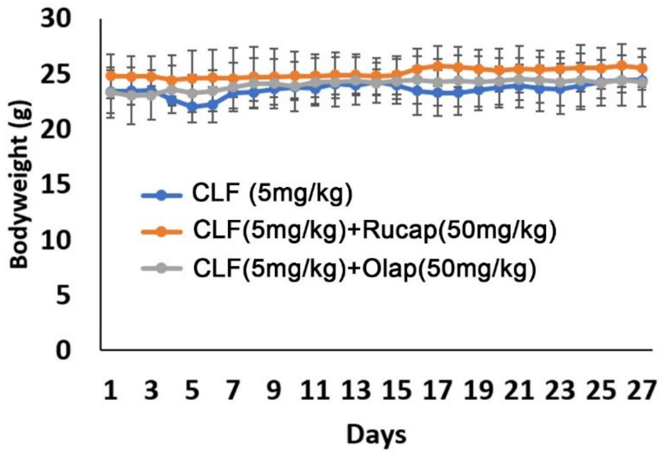


**Supplementary figure 11:** Tolerability assessed on non-tumor bearing female mice by administering CLF (5mg/kg IP 5d/w), CLF+ Rucaparib (50mg/kg oral gavage daily) and CLF+ Olaparib (50mg/kg oral gavage daily) for 4 weeks by body weight and BCS study. CLF and PARPis combinations showed no significant adverse effects.

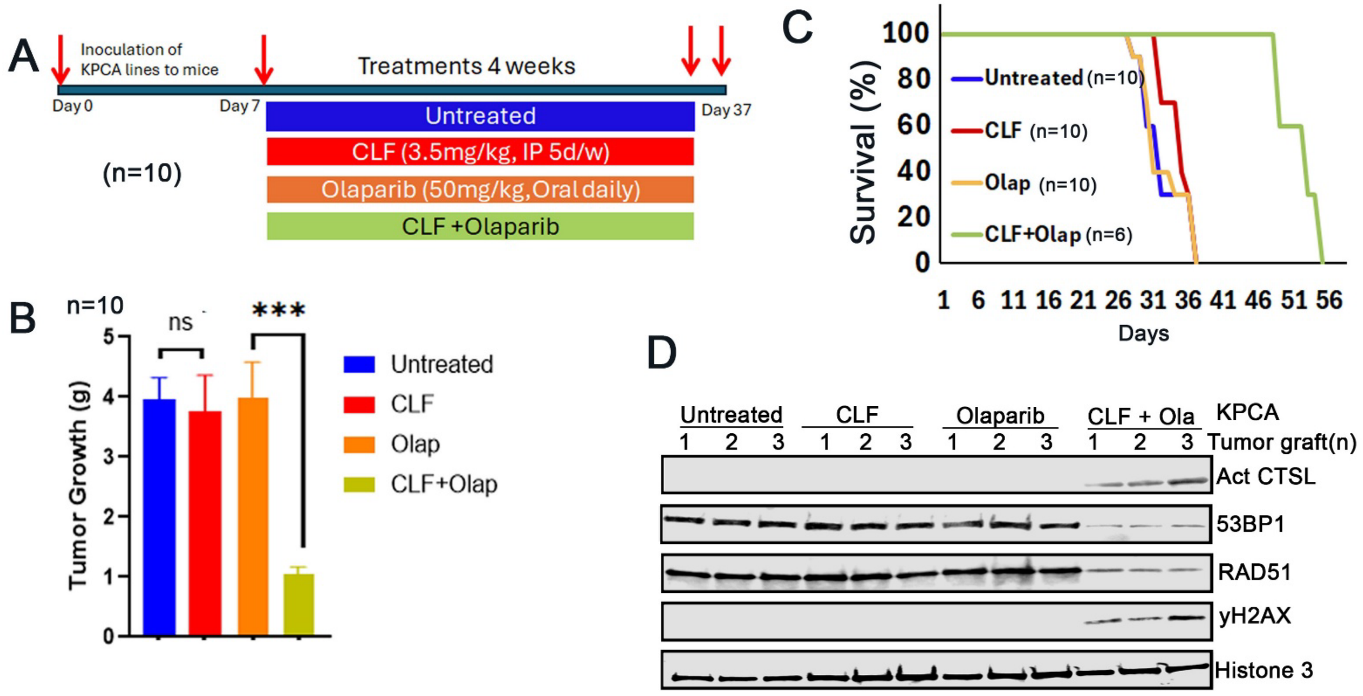


**Supplementary figure 12: Antitumor efficacy of CLF and olaparib in the KPCA syngeneic tumor model.**

Mice bearing KPCA tumors were treated with vehicle control, CLF, olaparib, or the CLF/olaparib combination for 4 weeks, and tumor growth and survival were monitored. (A) Treatment schedule. (B) Tumor growth was inhibited by CLF and olaparib, with greater antitumor efficacy observed with the combination. (C) Kaplan–Meier survival analysis with pairwise log-rank tests showed that CLF plus olaparib significantly improved survival compared with vehicle control or either monotherapy (**$p < 0.01$). (D) Western blot analysis of nuclear fractions from KPCA tumor tissues was performed to assess nuclear CTSL, 53BP1, RAD51 and the nuclear control Histone H3 levels.

**Supplementary Table 1**

**Sources of Materials Used**

| Sl no | Name of the chemicals/reagent/plasmid | Catalog number | RRIDs | Manufacturer |
|---|---|---|---|---|
| 1 | Clofarabine | HY-A0005/CS-0373 | | MedChemExpress (Monmouth Jct., NJ) |
| 2 | Importazole | HY-101091/CS-6189 | | |
| 3 | Eltanexor (KPT-8602) | S8397 | | Selleckchem (Houston, TX) |
| 4 | Niraparib | MK4827 | | Chemietek (Indianapolis, IN) |
| 5 | Veliparib | CT-A888 | | |
| 6 | Rucaparib | - | | Clovis Oncology (Boulder, CO) |
| 7 | Cytarabine hydrochloride, fludarabine and gemcitabine hydrochloride | NCI- DCTD | | Approved Oncology Drugs-NCI- DCTD |
| 8 | Z-FY(tBU)-DMK | sc-222423 | | Santa Cruz Biotechnology (Dallas, TX) |
| 9 | Olaparib | sc-302017A | | |
| 10 | Doxorubicin | sc-280681 | | |
| 11 | Z-DEVD-FMK | #14414 | | Cayman Chemical (Ann Arbor, MI) |
| 12 | CA-074Me | #18469 | | |
| **Sl no** | **Name of the Antibodies/Plasmid/Kits** | **Catalog number** | **RRIDs** | **Manufacturer** |
| 1 | Anti-CRM1 | sc-74454 | AB_1122704 | Santa Cruz Biotechnology (Dallas, TX) |
| 2 | anti- PAX8 | sc-81353 | AB_1127048 | |
| 3 | anti-GAPDH | sc-47724 | AB_627678 | |
| 4 | anti-C9 tag | sc-390000 | AB_2894830 | |
| 5 | anti-KPNB1 | sc-137016 | AB_2133993 | |
| 6 | anti- KPNA1 | sc-101292 | AB_2133664 | |
| 7 | anti-β actin | sc-47778 | AB_626632 | |
| 8 | anti- α tubulin | sc-5286 | AB_628411 | |
| 9 | Anti-Snai1/ Snail1 | sc-271977 | AB_10709902 | |

| | | | | |
|---|---|---|---|---|
| 10 | Anti-CRM1 | #46249S | AB_2799298 | |
| 11 | anti- PARP | #9541S | AB_331426 | |
| 12 | Anti-cleaved caspase-3 | #9661 | AB_2341188 | Cell Signaling Technology (Danvers, MA) |
| 13 | anti-53BP1 | #11940 | AB_2637071 | |
| 14 | anti- Flag tag | #9291S | AB_10950495 | |
| 15 | cell lysis buffer | #9803S | | |
| 16 | anti- RAD51 | #ab133534 | AB_2722613 | Abcam |
| 17 | Anti-CTSL | #10486-R221 | SCR_003697 | Sino Biological |
| | IRDye 680 LT goat anti mouse IgG, red | 926-68020 | AB_2687826 | |
| 17 | IRDye 800CW Donkey anti-Mouse IgG, green | 926-32212 | AB_2716622 | Li-Cor, Lincoln, NE, U.S.A. |
| 18 | IRDye 800 goat anti rabbit IgG, green, | 926-32211 | AB_621843 | |
| | IRDye 680RD Donkey anti-Rabbit IgG, red | 926-68073 | AB_10954442 | |
| 19 | Pacific Blue™ Annexin V | #11 858 777 | | Roche, Basel, Switzerland |
| 20 | shCTSL -TRC1.5 plasmid | SHCLNG-NM_001912 | | Sigma Aldrich |
| 21 | anti- RAD51 | #ab133534 | AB_2722613 | Abcam |
| 22 | pcDNA3.1-hCathepsin L construct | #11250 | Addgene_11250 | Addgene |
| 23 | recombinant human CTSL (rCTSL) protein | 952-CY | | R&D Systems |
| 24 | recombinant human CTSB (rCTSB) protein | 10483-H08H | | Sino Biological |
| 25 | NE-PER nuclear and CE-PER | #78833 | | Life Technologies |
| 26 | anti-hFAP | # AF3715 | AB_2102369 | R & D systems |
| 27 | Magic Red® Cathepsin L Assay | #942 | | ImmunoChemistry Technologies LLC |

**Supplementary Table 2**

**Cell lines, primary tumors and PDX cells**

| **Cell lines/RRIDs** | **Media** | **Source** |
|---|---|---|
| OVCAR-3 (CVCL_0465),<br>OVCAR-4 ( CVCL_1627) ,<br>OVCAR-5( CVCL_1628),<br>OVCAR-8( CVCL_1629),<br>HeyA8 (CVCL_8878),<br>A549(CVCL_A459),<br>H460 (CVCL_H460),<br>MDA-MB-231 CVCL_0062)<br>MDA-MB-436 (CVCL_0623) | RPMI (10% FBS+ 1% Penicillin/streptomycin) | American Type Culture Collection (ATCC), Manassas, Virginia, (SCR_001672) |
| H1048( CVCL_1453),<br>H1882( CVCL_1504),<br>KLE( CVCL_1329),<br>RL95-2 ( CVCL_0505) and<br>Hec1B( CVCL_0294) | DMEM/F12 (10% FBS+ 1% Penicillin/streptomycin) | |
| HeyA8-MDR ( CVCL_8879) | RPMI (10% FBS+ 1% Penicillin/streptomycin) | |
| COV362(CVCL_2420) | RPMI (10% FBS+ 1% Penicillin/streptomycin) | Dr. Kaufmann lab, Mayo Clinic, Rochester MN |
| PEO1(CVCL_2686) and PEO/OLTRC4 | RPMI (10% FBS+ 1x Penicillin/streptomycin) | Xinyan Wu lab, Mayo Clinic, Rochester MN |
| PDXs- PH610, PH747, PH077R, PH242, PH081, PH235, PH115, PH088 and PH442<br>Primary tumor- OV1182 and OV1183 | DMEM/F12 (15% FBS+ 1% Penicillin/streptomycin + 1x L-glutamine) | S. John Weroha lab, Mayo Clinic, Rochester MN |
| Patient derived Ovarian cancer Ascitic cells (OVAs) | DMEM/F12 (15% FBS+ 1% Penicillin/streptomycin + 1x L-glutamine) | University of Minnesota Cancer Center Tissue Procurement Facility and Mayo Clinic Rochester MN |
| HCC-1395, HCC-1937, MDA-MB-453 | RPMI (10% FBS+ 1% Penicillin/streptomycin) | Kuntian Lu, Mayo Clinic, Rochester MN |
| SNU539 and SNU1077 | DMEM/F12 (10% FBS+ 1% Penicillin/streptomycin) | Ja-Lok Ku, Cancer Research Institute,<br>Seoul, Korea |
| ARK1-UMMT | DMEM/F12 (10% FBS+ 1% Penicillin/streptomycin) | Alessandro D. Santin, Yale University, New Haven, CT |

**Supplementary Table 3 : Primary Ovarian Cancer Ascites (OVA), PDX (PH) and primary tumor (OV)samples**

| OVA | **Histology** | **Stage** | **Grade** | **Germline BRCA Mutations** | **Somatic BRCA and TP53 Mutations** |
|---|---|---|---|---|---|
| OVA-1 | High Grade Serous | 3C | 3 | Not tested | TP53- R273C |
| OVA-2 | High Grade Serous | 3C | 3 | Negative | TP53-P318fs*27 |
| OVA-9 | High Grade Serous | 3C | 3 | Negative | BRCA2 c.5415_5418del (p.Asn1805Lysfs*9) |
| OVA-10 | High Grade Serous | High | Unknown | Not tested | Not tested |
| OVA-11 | High Grade Serous | High | 3 | Not tested | Not tested |
| OVA-12 | High Grade Serous | 3C | 3 | Negative | TP53-R273C |
| OVA-15 | High Grade Serous | IVA | 2 | Negative | TP53-C124fs*46 |
| OVA-16 | High Grade Serous | 3A | 3 | Negative | Negative |
| OVA-22 | High Grade Serous | Unknow n | Unknown | Not tested | TP53 Deleterious- E258fs* |
| OVA-28 | High Grade Serous | IIIC | high | Negative | Not tested |
| OVA-29 | High Grade Serous | IVA | high | Negative | TP53- R248Q |
| OVA-30 | | III | | Not tested | Not tested |
| OVA-33 | High Grade Serous | IVB | 3 (high) | Negative | TP53- p.E339* |
| OVA-34 | High Grade Serous | IIIC | Low | Negative | Negative |
| PH610 | High Grade Serous | IV | high | BRCA WT | Not tested |
| PH747 | High Grade Serous | IIIC | 3 | Negative | TP53- p.R175H c.524G>A BRCA1- c.2138C>G (p.Ser713Ter) |
| OV1182 | High Grade Serous | IIIC | high | Negative | TP53- splice site 993+1G>A |
| OV1183 | High Grade Serous | IVB | high | BRCA1- c.2359dup (p.Glu787Glu787Glyfs*3 ) | Not tested |

**Supplementary Table 4**: List of ovarian cancer cells categorized based on drug induced nCTSL trafficking

| **Cancer types** | **Name of cell lines** | **CLF-r** | **CLF-nr** | **CLF resistant** |
|---|---|---|---|---|
| | | ***Presence of nCTSL-*** **CLF Monotherapy** | ***Presence of nCTSL*** **CLF +Olaparib** | ***Presence of nCTSL*** **CLF +KPT8602** Or **Olaparib+KPT8602** |
| Ovarian Cancer | PEO1 | Yes | - | |
| | PEO1/OLTRC4 | - | Yes | |
| | PEO1/ABTr#3 | - | Yes | |
| | HeyA8 | Yes | - | |
| | HeyA8-MDR | - | Yes | |
| | OVCAR-8 | - | Yes | |
| | OVCAR-3 | - | Yes | |
| | OVCAR-4 | - | Yes | |
| | OVCAR5 | Yes | - | |
| | COV362 | Yes | - | |
| | Primary tumor-PH1182 | Yes | - | |
| | Primary tumor-PH1183 | - | Yes | |
| | OC-Ascitic cells-OVA-2 | Yes | - | |
| | OC-Ascitic cells-OVA-11 | Yes | | |
| | OC-Ascitic cells-OVA-12 | - | Yes | |
| | PDX-PH610 | Yes | - | |
| | PDX-PH077R | Yes | - | |
| | PDX-PH242 | Yes | - | |
| | PDX- PH747 | - | Yes | |
| | PDX- PH081 | - | Yes | |
| | PDX-PH235 | - | Yes | |
| | PDX-PH115 | - | Yes | |
| | PDX-PH088 | - | Yes | |
| | PDX-PH442 | - | Yes | |

CLF-r: Responsive to CLF, CLF-nr; Responsive to CLF/Olaparib combination, CLF Resistant; Resistant to CLF/olaparib combination.

**Supplementary Table 5**: List of other cancer types categorized based on drug induced nCTSL trafficking

| Cancer types | Name of cell lines | CLF-r | CLF-nr | CLF resistant |
|---|---|---|---|---|
| | | ***Presence of nCTSL*- CLF Monotherapy** | ***Presence of nCTSL* CLF +Olaparib** | ***Presence of nCTSL* CLF +KPT8602** Or **Olaparib+KPT8602** |
| Breast Cancer | TNBC :MDA-MB-436 | Yes | - | |
| | TNBC: MDA-MB-231 | Yes | - | |
| | TNBC:HCC1395 | Yes | - | |
| | TNBC:HCC1937 | Yes | - | |
| | TNBC: MDA-MB-453 | - | - | Not screened |
| | Luminal: MCF-7 | - | Yes | |
| Cervical Cancer | OV2008 | Yes | - | |
| | OV2008-C13 | - | Yes | |
| Endometrial Cancer | KLE | - | Yes | |
| | ARK1-UMMT | Yes | - | |
| | RL95-2 | - | - | Yes (CLF-r) |
| | Hec1B | - | - | Not screened |
| Uterine Carcinosarcoma | SNU539 | Yes | - | |
| | SNU1077 | - | Yes | |
| Non Small Cell Lung Cancer | A549 | - | Yes | |
| | H460 | - | Yes | |
| Small Cell Lung Cancer | H1048 | - | - | Not screened |
| | H1882 | - | - | Not screened |

CLF-r: Responsive to CLF, CLF-nr; Responsive to CLF/Olaparib combination, CLF resistant; Resistant to CLF/olaparib combination.
